# Task Engagement Enhances Population Encoding of Stimulus Meaning in Primary Auditory Cortex

**DOI:** 10.1101/240556

**Authors:** Sophie Bagur, Martin Averseng, Diego Elgueda, Stephen David, Jonathan Fritz, Pingbo Yin, Shihab Shamma, Yves Boubenec, Srdjan Ostojic

## Abstract

The main functions of primary sensory cortical areas are classically considered to be the extraction and representation of stimulus features. In contrast, higher cortical sensory association areas are thought to be responsible for combining these sensory representations with internal motivations and learnt associations. These regions generate appropriate neural responses that are maintained until a motor command is executed. Within this framework, responses of the primary sensory areas during task performance are expected to carry less information about the behavioral meaning of the stimulus than higher sensory, association, motor and frontal cortices. Here we demonstrate instead that the neuronal population responses in the early primary auditory cortex (A1) display many aspects of responses generally associated with higher-level areas. A1 activity was recorded in awake ferrets while they were either passively listening or actively discriminating two periodic click trains of different rates in a Go/No-Go paradigm. By applying population-level dimensionality reduction techniques, we found that task-engagement induced a shift in the nature of the encoding from a sensory-driven representation of the two stimuli to a behaviorally relevant representation of the two categories that specifically enhances the target stimulus. We demonstrate that this shift in encoding relies partly on a novel mechanism of change in spontaneous activity patterns upon engagement in the task. We show that this population-level representation of stimuli in A1 population activity bears strong similarities to responses in the frontal cortex, but appears earlier following stimulus presentation. Analysis of neural activity recorded in various Go/No-Go tasks, with different sounds and reinforcement paradigms, reveals that this striking population-level enhancement of target representation is a general property of task engagement. These findings indicate that primary sensory cortices play a highly flexible role in the processing of incoming stimuli and implement a crucial change in the structure of population activity in order to extract task-relevant information during behavior.

## Introduction

How and where in the brain are sensory representations transformed into abstract percepts? Classical anatomical and physiological studies have suggested that this transformation occurs progressively along a cortical hierarchy. Primary sensory areas are commonly believed to process and extract high-level physical properties of stimuli, such as orientations of visual bars in the primary visual cortex or abstract sound features in the primary auditory cortex^1,2^. These fundamental sensory features are then integrated and interpreted as behaviorally meaningful sensory objects in sensory scenes, and relayed to higher cortical areas, which extract increasingly task-relevant abstract information. Prefrontal, parietal and premotor areas lie at the apex of the hierarchy^3,4^. They integrate inputs from different sensory modalities, transform sensory information into categorical percepts and decisions, and store them in working memory until the time when the appropriate motor action needs to be executed^5,6^.

According to this classical feedforward picture, primary sensory areas are often considered as playing a largely static role in extracting and encoding high-level stimulus physical attributes^7–10^. However a number of recent studies in awake, behaving animals have challenged this view, and shown that the information represented in primary areas in fact strongly depends on the behavioral state of the animal. Motor activity, arousal, learning and task-engagement have been found to strongly modulate responses in primary visual, somatosensory, and auditory cortices^11–25^. Effects of task-engagement have been particularly investigated in the auditory cortex, where it was found that receptive fields of primary auditory cortex neurons adapt rapidly to behavioral demands when animals engage in various types of auditory discrimination tasks^26–30^. These observations have been interpreted as signatures of highly flexible sensory representations in primary cortical areas, and they raise the possibility that these areas may be performing computations more complex than simple extraction and transmission of processed stimulus features to higher-order regions.

An important limitation of many previous studies^26–30^ is that they relied mostly on single-cell analyses, which characterized the selectivity of individual neurons to sensory stimuli. Here we show that simple population analyses reveal that task-engagement induces a shift in the primary auditory cortex from a sensory-driven representation to a representation of the behavioral meaning of stimuli, analogous to the one found in the frontal cortex. We first analyzed the responses during a temporal auditory discrimination task, in which ferrets had to distinguish between Go (Reference) and No-Go (Target) stimuli corresponding to click trains of different rates. The activity of the same neural population was recorded when the animals were engaged in the task, and when they passively listened to the same stimuli. Both single cell and population analyses showed that task-engagement decreased the accuracy of encoding the physical attributes of stimuli. Population, but not single-cell, analyses however revealed that task-engagement induced a shift towards an asymmetric representation of the two stimuli that enhanced target-evoked activity in the subspace of optimal decoding. This shift was in part enabled by a novel mechanism based on the change in the pattern of spontaneous activity during task engagement.

Performing identical analyses developed on this task to independent data sets collected in A1 during other behavioral discrimination tasks demonstrated that these findings can be well generalized, independently of the type of stimuli, behavioral paradigm or reward contingencies. Specifically, in all tasks, we found an enhanced representation of the target stimuli, defined as those stimuli that induced a change in the animal’s ongoing behavior. Furthermore, in tasks that displayed a shift in the spontaneous firing rates of neurons, this task-adaptive encoding was partly mediated by a re-patterning of the population spontaneous activity, offering a functional interpretation for this previously observed phenomena of task-evoked changes in spontaneous activity ^19^.

Finally, a comparison between population activity in A1 and single-cell recordings in the frontal cortex revealed strong similarities. However, the target-driven representation of behavioral meaning appeared in A1 very rapidly following stimulus presentation, hence it was unlikely to be solely due to immediate top-down influences from frontal cortex. Altogether, our results suggest that task-relevant, abstracted information is present in primary sensory cortices, and can be read out by neurons in higher order cortices.

## RESULTS

### Task engagement degrades the encoding of stimulus physical features in A1

We recorded the activity of 370 units in the primary auditory cortex (A1) of two awake ferrets in response to periodic click trains. The animals were trained using a conditioned avoidance paradigm^26^ to lick water from a spout during the presentation of a class of reference stimuli and to stop licking following a target stimulus (Animal 1: 83% hit +/−3% s.e.m; Animal 2: 69% hit +/−5% s.e.m) (Fig. 1a; see Methods). Target stimuli thus required a change in the ongoing behavioral output while reference stimuli did not. Each animal was trained to discriminate low vs high click rates, but the precise rates of reference and target click trains changed in every session. The category choice was opposite in the two animals to avoid confounding effects of stimulus rates (low/high) and behavioral category (reference/target). Thus, the target for one ferret was high click train rates, and the target for the other ferret was low click train rates. In each session, the activity of the same set of single units was recorded during active behavior (task-engaged condition) and during passive presentations of the same set of auditory stimuli before and after behavior (passive conditions).

**Fig 1.**
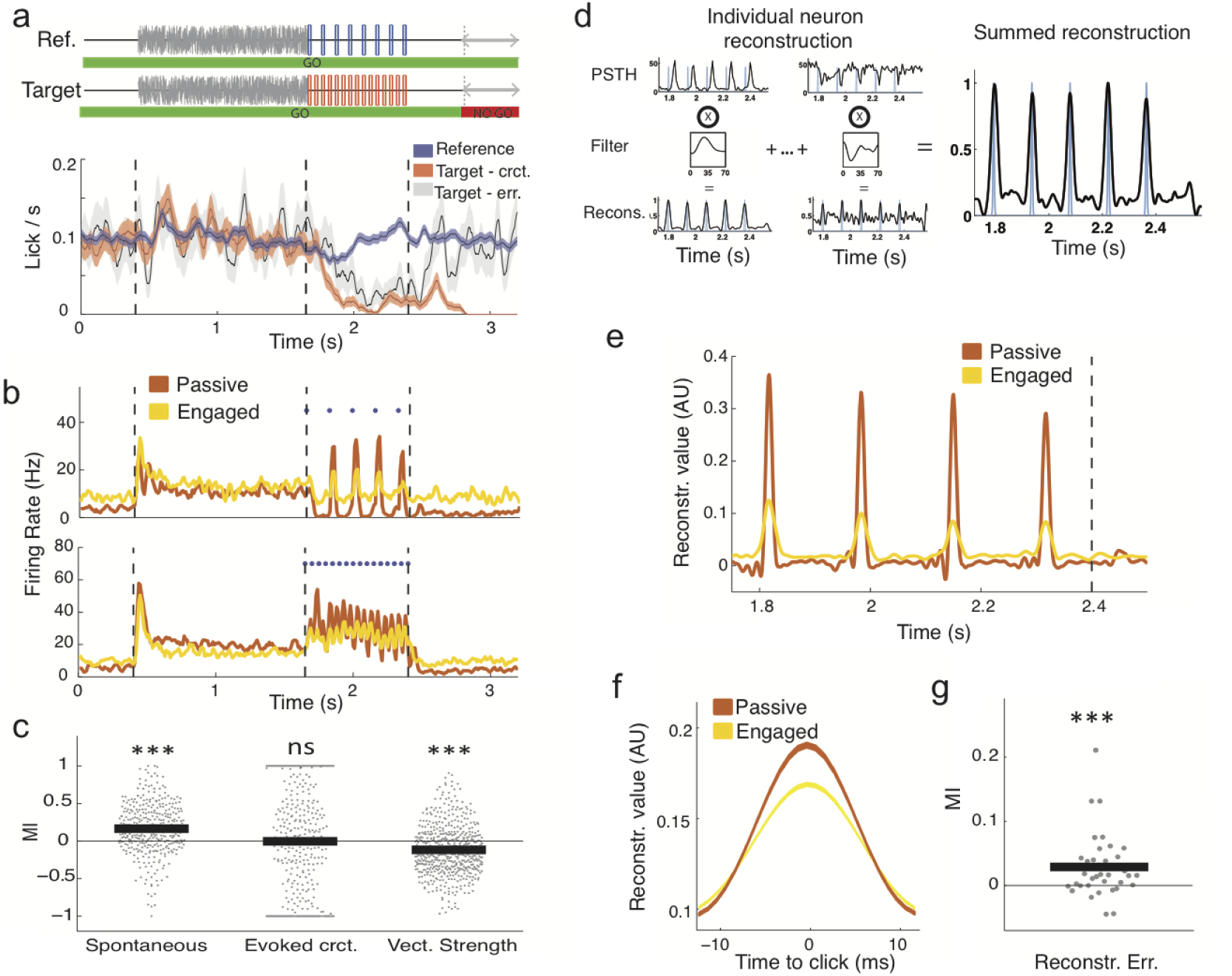
Task structure and neural encoding of click times in A1. *a. Structure of the click-train discrimination task and average behavior of the two animals. Each sound sequence is composed of 0.4s silence then a 1.25s long white noise burst followed by a 0.8s click train and a 0.8s silence. On each block the ferret is presented with a random number (1-7) of reference stimuli (top) preceding a target stimulus (bottom), except on catch trials with no target presentations. On blocks including a target, the animal had to refrain from licking during the final 0.4s of the trial, the no go period, to avoid a mild tail shock. (error bars are +/−sem)* *b. PSTH of two example units during reference sequences in the passive and engaged state. Note that in the task-engaged state, the units show enhanced firing during the initial silent period of spontaneous activity and reduced phase locking to the stimulus*. *c. Modulation index of each unit for spontaneous firing rate, spontaneous-corrected click-evoked firing rate and vector strength showing higher spontaneous firing rates and lower vector strength in the task-engaged state. The vector strength was only calculated for units firing above 1 Hz and values for both reference and target are shown. SEM error bars are not shown because not visible at this scale: 0.017, 0.037 and 0.013 respectively. (one-sample two-sided Wilcoxon signed rank test with mean 0, n=370, 574, 370, zval=−8.99, p=2.57e-19; zval=−0.07, p=0.94; zval=−8.82, p=1.16e-18; ***: p<0.001)*. *d. Schematic of stimulus reconstruction algorithm. Using PSTHs from half of the trials, a time-lagged filter is fitted to allow optimal reconstruction of the stimulus for each individual unit. Individual reconstructions are summed to obtain a population reconstruction (far right)*. *e. Stimulus reconstruction from an example session showing degraded reconstruction in the task-engaged state*. *f. Mean click reconstruction in passive and engaged states*. *g. Modulation index of each session for stimulus reconstruction error. SEM error bar is not shown because not visible at this scale: 0.0014. (one-sample two-sided Wilcoxon signed rank test with mean 0, n=36; zval=−3.4092, p=6.51e-4; ***: p<0.001)*.

We first examined how auditory cortex responses and stimulus encoding depended on the behavioral state of the animal. In agreement with previous studies^14,19^, spontaneous activity often increased in the task-engaged condition, while stimulus-evoked activity was often suppressed (Fig. 1b). To quantify the changes in activity over the population, we used a modulation index of mean firing-rates between passive and task-engaged conditions, estimated in different epochs (Fig. 1c; see Methods). Spontaneous activity before stimulus presentation increased in the engaged condition (n=370 units, P<0.0001), while baseline-corrected stimulus-evoked activity did not change overall (n=370 units, P=0.94). These changes in average activity suggested that the signal-to-noise (SNR) ratio between stimulus-evoked and spontaneous activity paradoxically decreased when the animals engaged in the task.

To quantify in a more refined manner the timing of neural responses with respect to click-times, we computed the vector strengths of individual unit responses, a standard measure of phase-locked activity evoked by click trains^12,31^. Vector strengths quantify the amount of entrainment of the neural response to the clicks, and range from 1 for responses fully locked to clicks to 0 for responses independent of click timing. A vast majority of neurons (Passive Ref/Targ: 80%, 81% and Active Ref/Targ: 84%, 81%) displayed statistically significant vector strengths in both conditions. However vector strength decreased in the engaged condition compared to the passive condition (Fig. 1c; n=574 (287 units, 2 sounds), P<0.0001), independently of the rate of the click train and the identity of the stimuli (Fig. S1). This reduction in stimulus-entrainment further suggested that task engagement degraded the encoding of click-times in A1.

The change in activity between passive and task-engaged conditions was heterogeneous across the neural population. While stimulus-entrainment was on average reduced in the engaged condition, a minority of neurons increased their responses. One possibility is that such changes reflect an increased sparseness of the neural code. Under this hypothesis, the stimuli are represented by smaller pools of neurons in the task-engaged condition, but in a more reliable manner. To address this possibility, we built optimal decoders that reconstructed click timings from the activity of all simultaneously recorded neurons, in a trial-by-trial manner (Fig. 1d, Methods). We found that the reconstruction accuracy decreased in the task-engaged condition compared to the passive condition (Fig. 1e-g), confirming that encoding of click-times decreased during behavior.

In summary, the fine physical features of the behaviorally relevant stimuli became less faithfully represented by A1 activity when the animals were engaged in this discrimination task.

### During sound presentation target and reference stimuli can be equally classified from A1 responses in passive and engaged conditions

In the task-engaged condition, the animals were required to determine whether the rate of each presented click train was high or low. They needed to make a categorical decision about the stimuli and correctly associate them with the required actions, before using that information to drive behavior. We therefore asked to what extent the two classes of stimuli could be discriminated based on population responses in A1, in the task-engaged and in the passive conditions.

We first compared the mean firing-rates evoked by target and reference click trains. While some units elevated their activity for the target stimulus (Fig. 2a, left), others preferred the reference (Fig. 2a, right). Over the whole population, mean firing rates were not significantly different for target vs reference stimuli (Fig. 2b) or for low vs high rate click trains (Fig. S2a). This observation held in both passive and task-engaged conditions. Discriminating between the stimuli was thus not possible on the basis of population-averaged firing rates (see Fig. S2b).

**Fig 2.**
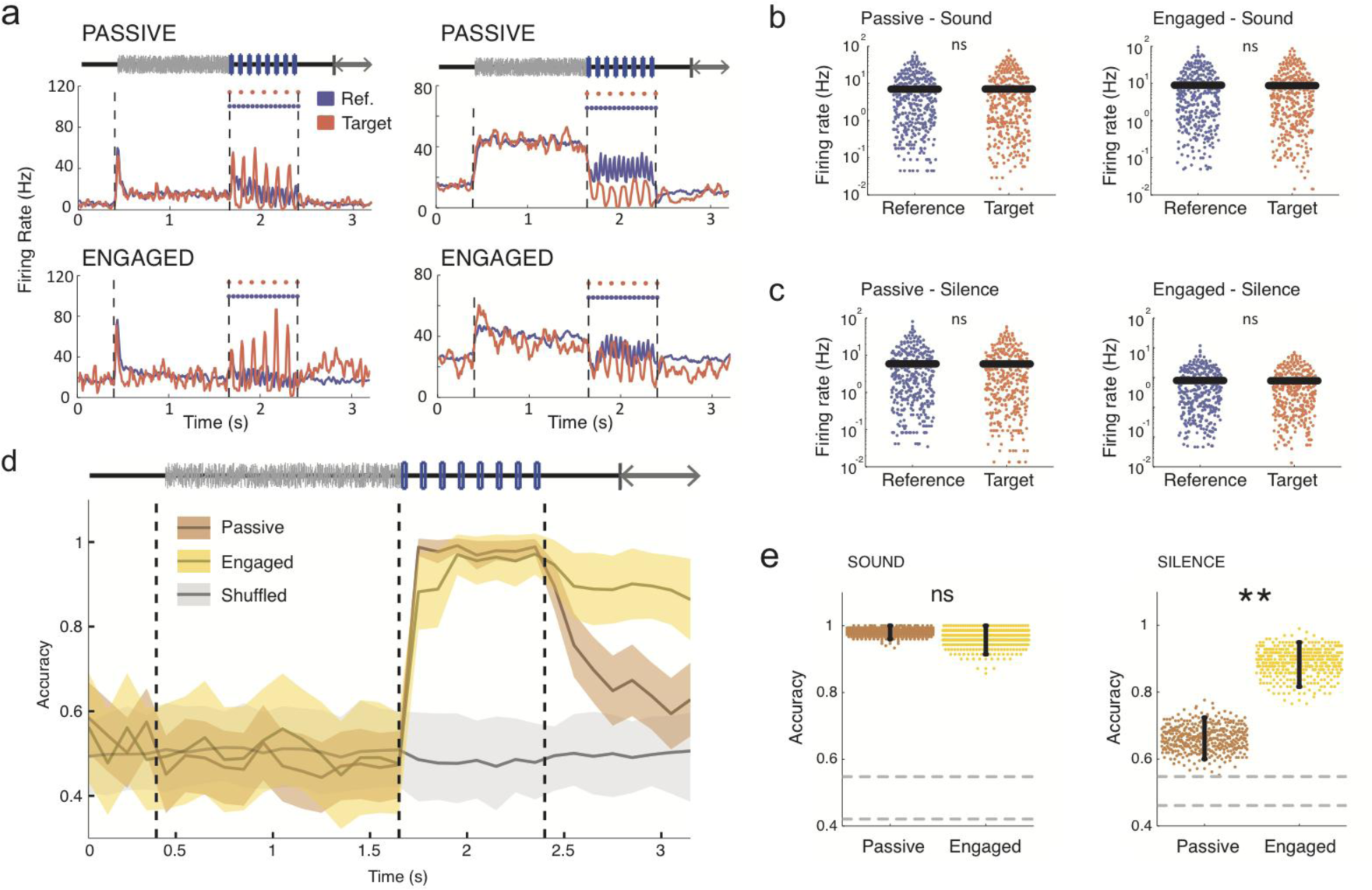
Discrimination of target and reference stimuli based on A1 activity. *a. PSTHs of two example units during reference (blue) and target (red) trials in the passive (top) and task-engaged (bottom) state. The unit on the left is target-preferring and the unit on the right is reference-preferring*. *b-c. Comparison of average firing rates on a log scale in passive (left) and engaged (right) between target and reference stimuli during the sound (b) and during the post-stimulus silence (c) periods. SEM error bars are not shown because not visible at this scale. (two-sided Wilcoxon signed rank, n=370; zval=0.34, p=0.73; zval=0.35, p=0.79; zval=−0.47, p=0.64; zval=−0.35, p=0.73)* *d. Accuracy of stimulus classification in passive and engaged states. In grey, chance level performance evaluated on label-shuffled trials. Error bars represent 1 std calculated over 400 cross-validations. e. Mean classifier accuracy during the sound (left) and silence period (right) in both conditions. Gray dotted lines give 95% confidence interval of shuffled trials. Error bars represent 95% confidence intervals. (n=400 cross validations; p=0.29 and p<0.0025; **: p<0.01)*

To take into account the heterogeneity of neural responses and quantify the ability of the whole population to discriminate between target and reference stimuli on an individual trial basis, we adopted a population-decoding approach. We used a simple, binary linear classifier that mimics a downstream readout neuron. The classifier takes as inputs the spike-counts of all the units in the recorded population, multiplies each input by a weight, and compares the sum to a threshold to determine whether a trial was a reference or a target. The weight of each unit was set based on the difference between the average spike-counts evoked by the two stimuli (Fig. S3 and Methods). This weight was therefore positive or negative depending on whether it preferred the target or reference stimulus. Different decoder weights were determined at every time-bin in the trial. The width of the time-bins (100ms) was larger than the inter-click intervals (Methods). Shorter time-bins increase the amount of noise but do not affect our main findings (Fig. S8A). Training and testing the classifier on separate trials allowed us to determine the cross-validated performance of the classifier, and therefore the ability to discriminate between the two stimulus classes based on single-trial activity in A1.

During stimulus presentation, the linear readout could discriminate target and reference stimuli with high accuracy in both passive and task-engaged conditions (Fig. 2d,e). Because the classifier performed at saturation during the sound epoch, it could be that differences between passive and active classifiers were masked by the substantial number of neurons provided to the classifiers. Decoders performing with lower numbers of neurons did not reveal any difference between the two behavioral states (Fig. S4a). Moreover this discrimination capability did not appear to be layer-dependent (Fig. S4b,c). The primary auditory cortex therefore appeared to robustly represent information about the stimulus class, independently of the decrease in the encoding of precise stimulus properties that occurs during task-engagement.

We next examined the discrimination performance during the silence immediately after stimulus offset. This silent period consisted of a 400ms interval followed by a response window, during which the animal learned to stop licking if the preceding stimulus was a target. As during the sound period, mean firing rates were not significantly different for the two types of stimuli during post-stimulus silence (Fig. 2c). Nevertheless, we found that discrimination performance between target and reference trials remained remarkably high throughout the post-stimulus silence in the task-engaged condition. In the passive condition, the decoding performance decayed during post-stimulus silence, but remained above chance level (Fig. 2d,e and Fig. S5b). The information about the stimulus class was thus maintained during the silent period in the neural activity in A1, but more strongly when the animal was actively engaged in the task. Moreover, a comparison between the decoders determined during the sound and after stimulus presentation showed that the encoding of information changed strongly between the two epochs of the trial (Fig. S6 and supplementary text).

### Task-engagement shifts encoding towards enhanced target-detection

We next examined in more detail the neural activity that underlies the classification performance in the two conditions. Target and reference stimuli play highly asymmetric roles in the Go/No-Go task design studied here as their behavioral meaning is totally different. As shown in Figure 1a, animals continuously licked throughout the task and only target stimuli elicited a change from this ongoing behavioral output while reference stimuli did not. We therefore sought to determine whether target- and reference-induced neural responses play similar or different roles in the discrimination between target and reference stimuli.

We first used dimensionality-reduction techniques to visualize the trajectories of the population activity in three dimensions (Fig. 3a, see Methods for details). The three principal dimensions were determined jointly for the passive and active data. This allowed us to visually inspect the difference in population dynamics and decoding axes between the two behavioral conditions. The average neural trajectories on reference and target trials strongly differ in the two behavioral conditions. In the passive condition, reference and target stimuli led to approximately symmetric trajectories around baseline spontaneous activity, suggesting that reference and target stimuli played essentially equivalent roles during the sound (Fig. 3a,c,d). In contrast, in the task-engaged condition, the activity evoked by reference and target stimuli became strongly asymmetric with respect to the decoding axes and the spontaneous activity (Fig. 3b,e,f).

**Fig 3.**
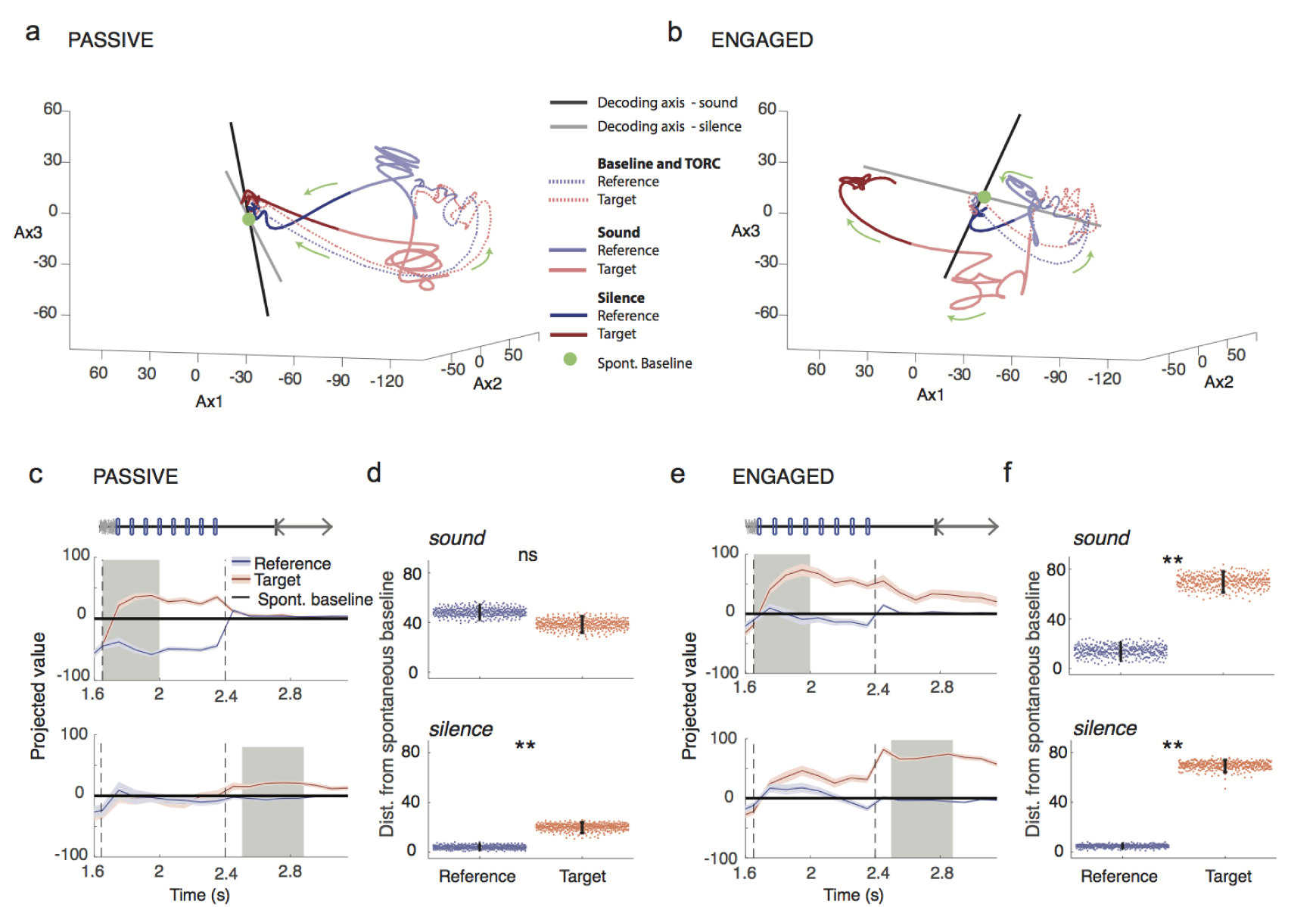
Task engagement induces shift from symmetric to asymmetric representation of target and reference stimuli. *a. Population response during target and reference stimuli in the passive state along the first three components identified using GPFA (see methods) on single trial data. The session begins at the baseline (green dot), followed by the TORC presentation, (dotted line) then the click presentation of either the target and the reference sound (light red and blue respectively) and finally to the post-sound silence period (dark red and blue). Note in particular that in the passive state, the reference and target activities move away symmetrically from the baseline point given by projection of spontaneous activity*. *b. As in a, for the task-engaged state. Note that in this state, target activity makes a much larger excursion from the baseline than reference activity. The axes are the same as in panel a, as the GPFA analysis was performed jointly on passive and engaged data*. *c. Projection onto the decoding axis of trial-averaged reference- and target-evoked responses for the whole neural population. A baseline value computed from pre-stimulus spontaneous activity was subtracted for each unit, so that the origin corresponds to the projection of spontaneous activity (shown by black line). Decoding axes determined during sound presentation and post-stimulus silence are respectively used for projections in the top and bottom rows. The periods used to construct the decoding axis are shaded in gray. Error bars represent 1 std calculated using decoding vectors from cross-validation. This procedure allows visualization of the distance between reference and target evoked projections (that corresponds to decoding strength) and the distance of the stimuli-evoked responses from the baseline of spontaneous activity can be interpreted as the contribution of each stimulus to decoding accuracy*. *d. Distance of reference and target projections from baseline in each condition during the sound and silence period. Error bars represent 95% confidence intervals (n=400 cross validations; p=0.15 & p<0.0025; **: p<0.01)*. *e. As in c for the engaged state*. *f. As in d for the engaged state. (n=400 cross validations; p<0.0025 & p<0.0025; **: p<0.01)*.

To further characterize the change in information representation between the two conditions, we examined the average inputs from target and reference stimuli to a hypothetical readout neuron corresponding to a previously determined linear classifier. This is equivalent to projecting the trial-averaged population activity onto the axis determined by the linear classifier, trained at a given time point in the trial. This procedure sums the neuronal responses after applying an optimal set of weights. It effectively reduces the population dynamics from N=370 dimensions (where each dimension represents the activity of an individual neuron) to a single, information-bearing dimension. The discrimination performance of the classifier is directly related to the distance between reference and target activity after projection, so that the projection allows us to visualize how the classifier extracts the stimulus category from the neuronal responses to the two respective stimuli. Projecting the spontaneous activity along the same axis provides moreover a baseline for comparing the changes in activity induced by the target and reference stimuli along the discrimination axis. As the encoding changes strongly between stimulus presentation and the subsequent silence (Fig. S6 and supplementary text), we examined two projections corresponding to the decoders determined during stimulus and during silence.

As suggested by the three-dimensional visualization, the projections on the decoding axes demonstrated a clear change in the nature of the encoding between the two behavioral conditions. In the passive condition, reference and target stimuli led to approximately symmetric changes around baseline spontaneous activity (Fig. 3c,d). In contrast, in the task-engaged condition, the activity evoked by reference and target stimuli became strongly asymmetric (Fig. 3e,f). In particular, the projection of reference-evoked activity remained remarkably close to spontaneous activity throughout the stimulus presentation and the subsequent silence in the task-engaged condition. The strong asymmetry in the engaged condition, and the alignment of reference-evoked activity were found irrespective of whether the projection was performed on decoders determined during stimulus (Fig. 3e,f, top) or during silence (Fig. 3e,f, bottom). The time-courses of the two projections were however different, with target-evoked responses rising very rapidly (Fig. 3e,f top) when projected along the first axis, but much more gradually when projected along the second axis (Fig. 3e,f, bottom). In both cases, however, our analysis showed that in the active condition the discrimination performance relies on an enhanced detection of the target.

The strong similarity between the projection of reference-evoked activity and the baseline formed by the projection of spontaneous activity is not due to the lack of responses to reference stimuli in the engaged condition. Reference stimuli do evoke strong responses above spontaneous activity in both passive and task-engaged conditions. However, in the task-engaged, but not in the passive condition, the population response pattern of the reference stimuli appears to become orthogonal to the axis of the readout unit during behavior. The strong asymmetry between reference- and target-evoked responses is therefore seen only along the decoding axis, but not if the responses are simply averaged over the population, or averaged after sign correction for the preference between target and reference (Fig. S7).

We verified that these results are robust across a range of time bins (10ms-200ms), allowing us to cover timescales both on the order of the click rate and much longer. Both the increase in post-sound decoding accuracy in the engaged state and the increased asymmetry of target/reference representation were observed at all time scales (Fig. S8a,b).

### Encoding of stimulus behavioral meaning in A1 is independent of motor activity and reflects behavioral outcomes

One simple explanation of the asymmetry between target- and reference-evoked responses could potentially be the motor-evoked neuronal discharge. Indeed, during task-engagment, the animals’ motor activity was different following target and reference stimuli as the animals refrained from licking before the No-Go window following the target stimulus but not the reference stimulus (Fig. 1a). As neural activity in A1 can be strongly modulated by motor activity^17^, such effects could potentially account for the observed differences between target- and reference-evoked population activity.

To assess the role played by motor activity in our findings, we first identified units with lick-related activity. To this end, we used decoding techniques to reconstruct lick timings from the population activity, and determined the units that significantly contributed to this reconstruction by progressively removing units until licking events could not anymore be detected from the population activity. We excluded a sufficient number of neurons (10%) such that a binary classifier using the remaining units could no longer classify lick and no-lick time points as compared with random data (p>0.4; Fig. 4a,b, see Methods). We then repeated the previous analyses after removing all of these units. The discrimination performance between target and reference trials remained high and significantly different between the passive and the task-engaged conditions during the post-stimulus silence (Fig. 4c,d), while projection of target- and reference-elicited activity on the updated decoders still showed a strong asymmetry in favor of the target (Fig. 4e,f). This indicated that the information about the behavioral meaning of stimuli was represented independently of any overt motor-related activity. In all subsequent analyses we excluded all lick-responsive neurons.

**Fig 4.**
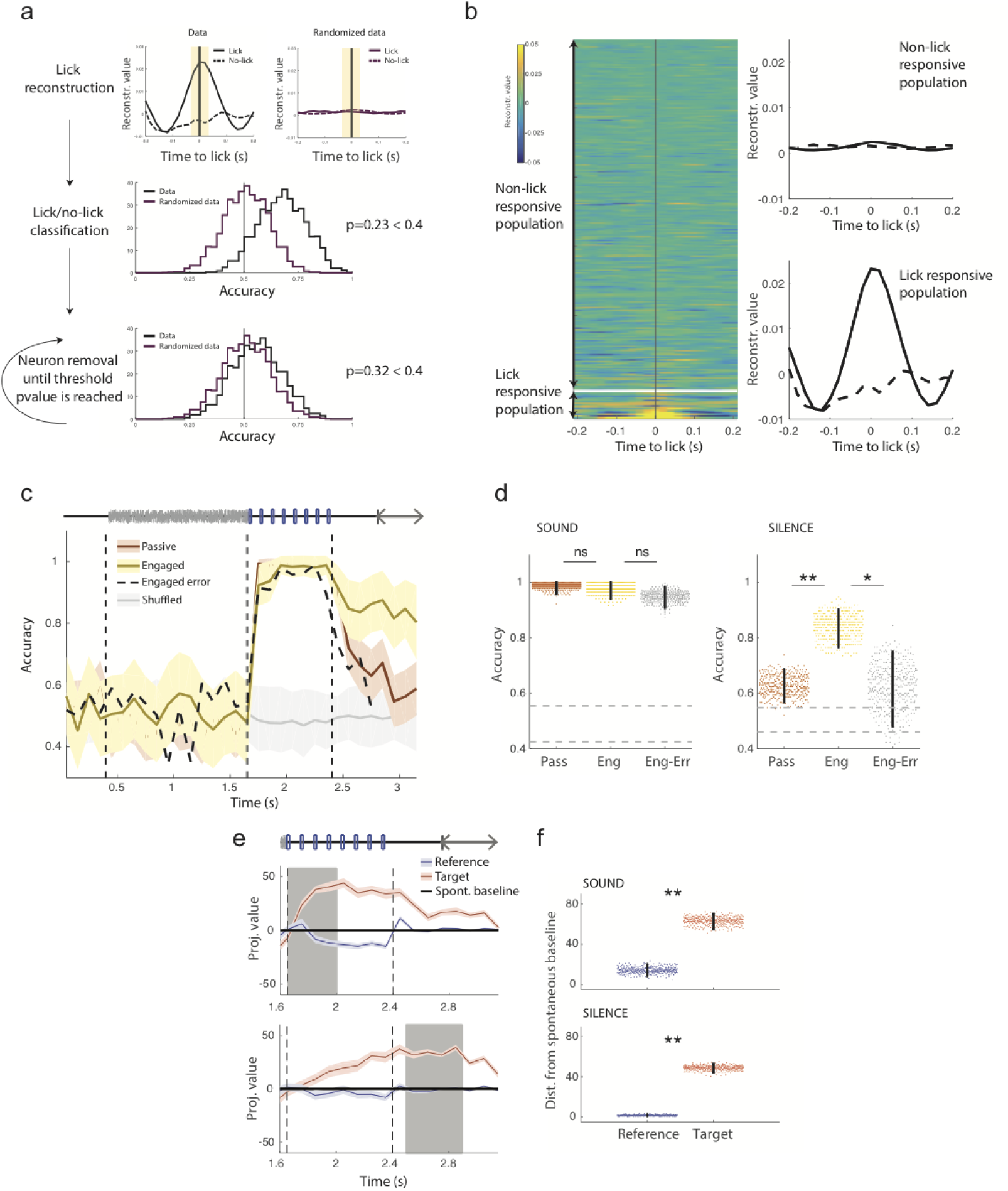
Relation between A1, motor activity and behavioural outcome. *a. Schematic of the approach used to identify lick responsive units to eliminate from population analysis. First, we reconstructed licks using optimal filters as with click reconstruction (Fig 1). To test whether this reconstruction allows to detect lick events, the filter is applied during licks and also during randomly selected time points with no licks (top left) to all units. Each event (lick or no-lick) can therefore be represented by a population vector constituted of the peak reconstruction values for all neurons. We evaluated the accuracy of classifying lick and no-lick time events using a linear decoder applied to this population vector (black distribution, middle panel). The same procedure was applied to randomized data (top right and purple distribution, middle panel) to test the significance of decoding and calculate a p-value (percentage of random data cross validations larger than real data cross validations). We then iteratively removed the best classification units (bottom plot) until the p-value was greater than 0.4 and the two distributions were indistinguishable. (see Methods for details)* *b. Results of reconstruction of lick events and removal of lick units. Left shows a heatmap of average lick reconstruction for all neurons ordered by their classification weight. Right shows the average reconstruction of lick and no-lick events using units retained for population analysis (non-lick responsive) and units excluded from the population analysis (lick-responsive)*. *c. Accuracy of stimulus classification in passive and engaged states using only non lick-responsive units. For the engaged state both correct and incorrect trials are shown. Note that after removal of lick-responsive units, the discrimination during post-stimulus silence is still enhanced in the task-engaged state on correct trials but is low during error trials. Error bars represent 1 std calculated over 400 cross-validations*. *d. Comparison of mean accuracy on passive, task-engaged correct and task-engaged error trials, during the sound (left) and post-stimulus silence periods (right). Error bars represent 95% confidence intervals. (n=400 cross validations; sound: pass/eng p=0.22, eng/err: p=0.87; silence: pass/eng p<0.0025, eng/err: p=0.012; *: p<0.05, **: p<0.01)* *e. Projection onto the decoding axis of baseline-subtracted population vectors during the engaged condition constructed using activity of non-lick responsive units only for the reference and target stimuli. Projections are shown onto the decoding axes obtained on early sound (top) and silence periods (bottom). The periods used to construct the decoding axis are shaded in gray. A baseline value computed from pre-stimulus spontaneous activity was subtracted for each unit, so that the origin corresponds to the projection of spontaneous activity (shown by black line). Error bars represent 1 std calculated using decoding vectors from cross-validation*. *f. Distance of reference and target projections from baseline in the engaged condition during the sound and silence periods. Error bars represent 95% confidence intervals (n=400 cross validations; p<0.0025 & p<0.0025; **: p<0.01)*.

Although the information present in A1 during the post-stimulus silent period could not be explained by motor activity, it appeared to be directly related to the behavioral performance of the animal. To show this, we classified population activity on error trials, in which the animal incorrectly licked on target stimuli, using classifiers trained on correct trials. Error trials showed only a slight impairment of accuracy during the sound presentation, but strikingly, the discrimination accuracy of the classifier during the post-stimulus silence on these trials dropped down to the performance level measured during passive sessions (Fig. 4c,e). This analysis therefore demonstrated a clear correlation between the behavioral performance and the information on stimulus category present during the silent period in A1.

Another aspect of neural activity that can be expected to change with task engagement is correlations between pairs of neurons. Our analysis so far has focused on the structure of population responses to external stimuli (signal correlations) but pairs of neurons display trial-to-trial fluctuations in activity (noise correlations) that can affect the population ability to encode information^32,33^. We found that task engagement decreased noise correlations on average (Fig. 5a,b; Fig. S9a), compatible with previous observations that attention reduces noise correlations^34^. Across the population, the range of changes was however very broad. To determine the influence of noise correlations on the population level, we repeated our analysis on simultaneously recorded data, using a modified linear decoder that takes noise correlations into account (the Fisher discriminant, see Methods). Our main findings appeared not to be sensitive to noise correlations. We were able to decode with high accuracy stimulus identity in passive and engaged states and observed an increase of stimulus memory in the engaged state as before (Fig. 5c). Projection onto this adjusted decoding axis showed a similar enhanced target representation in the engaged state, with the reference response lying along the projected baseline activity (Fig. 5d,e). Projection of responses using the linear classifier with and without taking noise correlations into account are strikingly similar across a range of timebins (Fig. S9b,c). Finally, a finer examination of the change between passive and engaged conditions showed that, contrary to previous observations^35^, noise correlations were most strongly reduced for pairs of neurons with opposite stimulus preference in our data set (Fig. S10b,c), which is expected to impair decoding of information (Fig. S10a).

**Fig 5.**
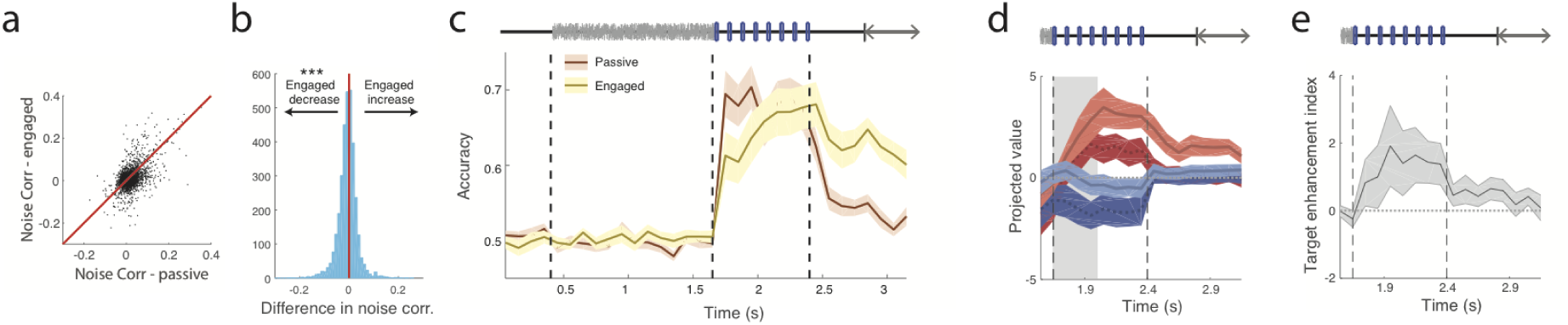
Task-induced changes in stimulus representation are independent of changes in noise correlations. *a. Comparison of noise correlations between pairs of neurons in the passive and engaged state. Red line indicates identity line*. *b. Histogram of correlation changes between the engaged and passive states showing a shift to lower values in the engaged state despite highly heterogeneous behavior across the population. (two-sided Wilcoxon signed rank, n=3361 pairs; zval=10.33, p=4.9E-25, ***:p<0.001)* *c. Accuracy of stimulus classification in passive and engaged states using simultaneously recorded, non lick-responsive units and applying a decoding vector corrected for noise correlations. Note that the increase in decoding accuracy during the silent period in the engaged state is still clearly visible. Error bars represent s.e.m over n=15 sessions*. *d. Projection onto the decoding axis determined during the sound period of trial-averaged reference (blue) and target (red) activity during the passive (dark colors) and the active (light colors) sessions. A baseline value computed from pre-stimulus spontaneous activity was subtracted for each neuron, so that the origin corresponds to the projection of spontaneous activity (shown by black line). Note that the target-driven activity lies further from the baseline in the active state and the reference-driven activity lies closer to baseline. The period used to construct the decoding axis is shaded in gray. Error bars represent s.e.m over n=15 sessions* e. Index of target enhancement induced by task engagement based on projections using the decoding axis determined during the sound. This value is positive if projected target activity is enhanced in the active state and projected reference activity is reduced. Error bars represent s.e.m over n=15 sessions.

### Mechanisms underlying the asymmetric, target-driven encoding during task-engagement

The previous analyses of population activity have shown that task engagement induces an asymmetric encoding, in which the activity elicited by reference stimuli becomes similar to spontaneous background activity when seen through the decoder. Two different mechanisms can potentially contribute to this shift between passive and engaged conditions: (i) the spontaneous activity changes between the two behavioral states such that its projection on the decoding axis becomes more similar to reference-evoked activity; (ii) stimulus-evoked activity changes between the states, inducing a change in the decoding axis and in the projections. In general, both mechanisms can be expected to contribute and their effects can be separated during different epochs of the trial.

To disentangle the effects of the two mechanisms, we chose a fixed decoding axis, and projected on the same axis the stimulus-evoked activity from both passive and engaged conditions. We then compared the resulting projections with projections of both passive and engaged spontaneous activity. We performed this procedure separately for decoding axes determined during sound and silence epochs.

Figure 6a (top) illustrates the projections along the decoding axis determined during the sound epoch in the engaged condition. Comparing the passive responses with the passive and engaged spontaneous activity revealed that the projection of passive reference-evoked activity was aligned during sound presentation with the projection of engaged, but not passive spontaneous activity (Fig. 6a top left). A similar observation held for the engaged responses throughout the sound presentation epoch (Fig. 6a top right). These projections remained similar regardless of whether the decoding axes were determined during the passive or the engaged conditions, as these two axes largely share the same orientation (Fig. S6e). Altogether, these results indicate that the change in spontaneous baseline activity during task engagement is sufficient to explain the strongly asymmetric, target-driven response observed early in the trial during sound presentation (Fig. 6b top).

**Fig 6.**
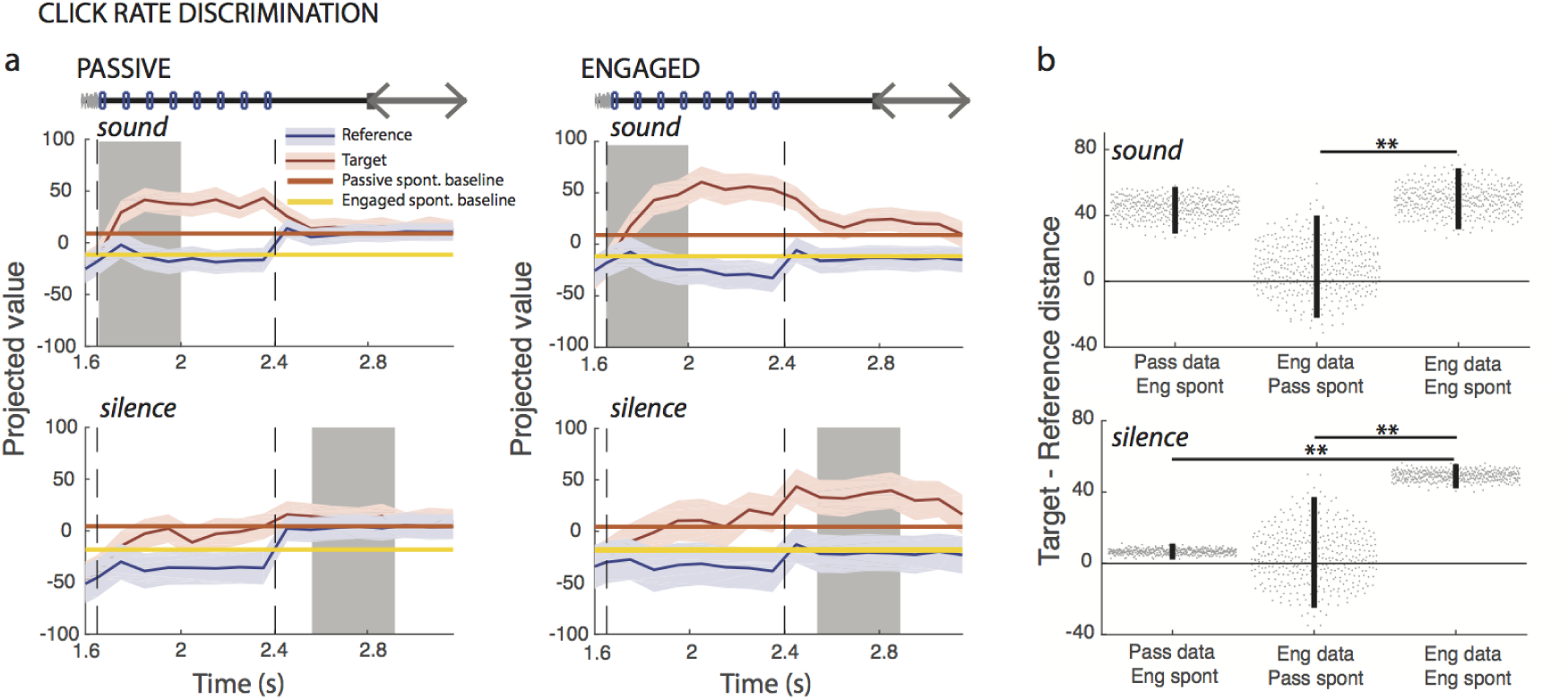
Shift in spontaneous activity contributes to change in asymmetry. Note that all analysis in this figure is done after excluding lick-responsive units in A1 as described in Fig 4. a. Projection onto the engaged decoding axis of reference- and target-evoked activity in the passive (left column) and engaged state (right column). Decoding axes determined during sound presentation and post-stimulus silence are respectively used for projections in the top and bottom rows. This figure differs from Fig 3c in which the spontaneous activity is subtracted before projection. Passive and engaged spontaneous activities after projection are shown by continuous lines. Error bars represent 1 std calculated using decoding vectors from cross-validation (n=400). b. Comparison of reference/target asymmetry for evoked responses in different states compared to different baselines given by passive or engaged spontaneous activity. Reference/target asymmetry is the difference of the distance of reference and target projected data to a given baseline. We examine three cases: (i) passive evoked responses, distances calculated relative to engaged spontaneous activity; (ii) engaged evoked responses, distances calculated relative to passive spontaneous activity; (iii) engaged evoked responses, distances calculated relative to engaged spontaneous activity. These values are shown during the sound (top) and the silence (bottom). In all three cases, the engaged decoding axis was used for projections. Decoding axes determined during sound presentation and post-stimulus silence are respectively used for projections in the top and bottom rows. Error bars represent 95% confidence intervals (n=400 cross validations; sound: p(col1,col3)=0.29 & p(col2,col3)<0.0025; silence: p(col1,col3)<0.0025 & p(col2,col3)<0.0025; **: p<0.01).

However, we reached a different conclusion when we examined the activity during the post-stimulus silence (Fig. 6a bottom). Repeating the same procedure as above, but projecting on the decoding axis determined during the post-stimulus silence revealed that the shift in spontaneous activity alone was not able to account for the asymmetry of the projected responses during the post-stimulus silence (Fig. 6b bottom). The target-driven, asymmetrical projections observed during this trial epoch therefore relied in part on a change in stimulus-evoked responses.

All together, we found that the changes in baseline spontaneous activity induced by the task engagement are key in explaining the enhancement of the target-driven, asymmetric encoding during sound presentation. As described in the above, the encoding axis during sound presentation is not drastically affected by task engagement. Instead, it is the population spontaneous activity that aligns with the reference-elicited activity with respect to the decoding axis. This observation in particular provides an additional argument against the possibility that the appearance of an asymmetrical representation is due to the asymmetrical motor responses to the two stimuli. Rather, the asymmetry is geometrically explained by baseline changes that precede stimulus presentation, and reflects the behavioral state of the animal.

### Sustained, target-driven, and behaviorally-gated responses of single cells in frontal cortex parallel population encoding in A1

The pattern of activity resulting from projecting reference- and target-elicited A1 activity on the linear readout is strikingly similar to previously published activity recorded in the dorsolateral frontal cortex (dlFC) of behaving ferrets performing similar Go/No-Go tasks (tone detect and two-tone discrimination in^36^). We therefore compared in more detail A1 activity with activity recorded in dlFC during the same click-rate discrimination task. When the animal was engaged in the task, single units in dlFC encoded the behavioral meaning of the stimuli by responding only to target stimuli, but remaining silent for reference stimuli (Fig. 6a bottom panel). Target-induced responses were moreover observed well after the end of the stimulus presentation, allowing for a maintained representation of stimulus category. The strong asymmetry of single-unit responses in dlFC clearly resembles the activity extracted from the A1 population by the linear decoder (Fig. 3 and 4). This suggests that the target-selective responses in the dlFC that reflect the cognitive decision process could in part be thought of as a simple readout of information already present in the population code of A1.

To further examine the relationship between dlFC single-unit responses and population activity in A1, we next compared the time course of the projected target-elicited data in A1 (Fig. 3e) and the population-averaged target-elicited neuronal activity in dlFC (Fig. 7a bottom panel) during active sessions. As mentioned above, the optimal decoding axes for A1 activity changes between the stimulus presentation epoch and the silence that follows (Fig. S6). The time-course of the projected A1 activity depends strongly on the axis used for the projection. When projecting on the axis determined during stimulus presentation, the target-elicited response in A1 was extremely fast (0.08s +/− 0.009 std) compared to the much longer response latency in the population-averaged response of dlFC neurons (0.48s +/− 0.12 std) (Fig. 7b). In contrast, when projecting on the axis determined during post-stimulus silence, the target-elicited response in A1 was slower (0.21s +/− 0.03 std) and closer to the population-averaged response in the dlFC (note that a fraction of individual units in dlFC display a very fast responses not reflected in the population average, see Fritz et al. 2010). Our analyses therefore identified two contributions to target-driven population dynamics in A1, a fast component absent in population-averaged dlFC activity and a slower component similar to population-averaged activity in dlFC, thus pointing to a possible contribution of an A1-FC loop that could be engaged during auditory behavior.

**Fig 7.**
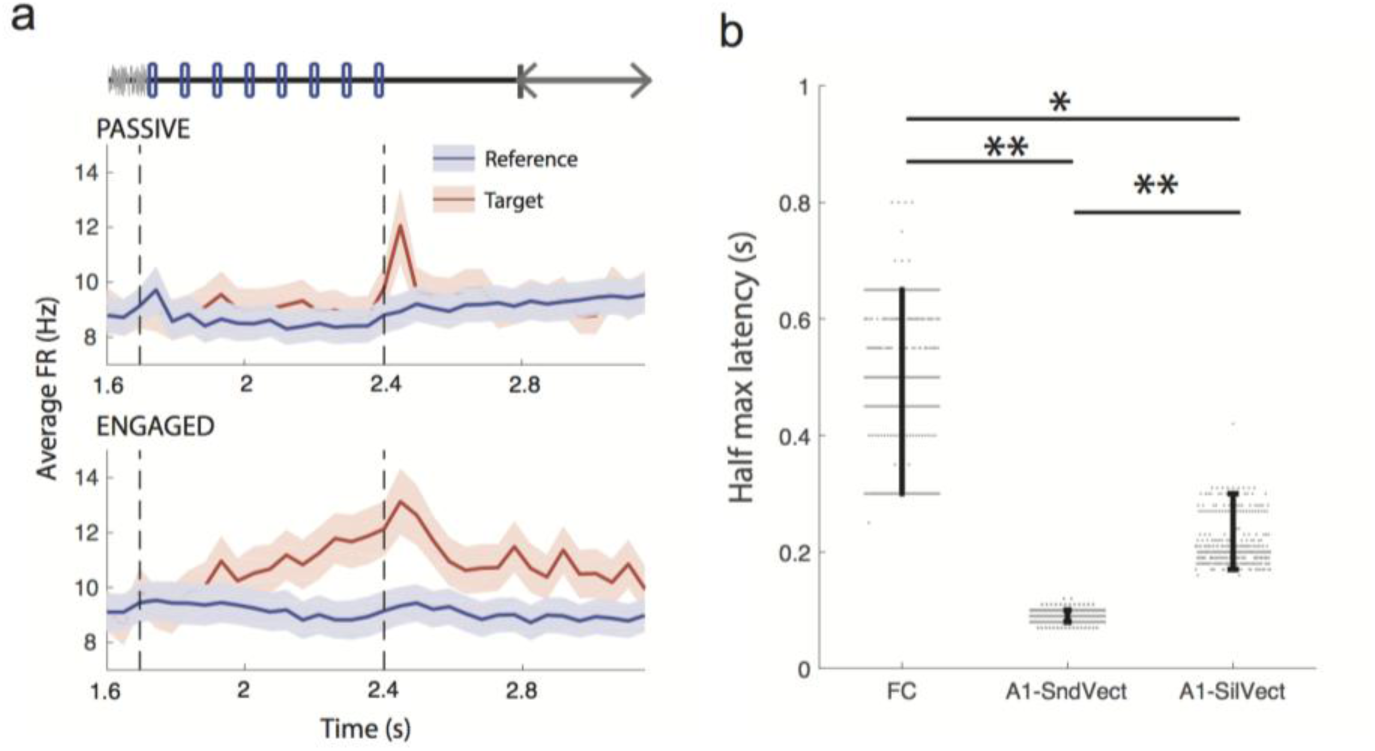
Persistent, asymmetric response to target and reference stimuli in frontal cortex. Note that all analysis in this figure is done after excluding lick-responsive units in A1 as described in Fig 4. a. Average PSTHs of all frontal cortex units in response to target and reference stimuli in both passive and engaged conditions. Note that the response to the target in the task-engaged state is very clear and appears late during the sound. Error bars: s.e.m over all units (n=102) b. Latency to half-maximum response for frontal cortex (for average PSTHs) and primary auditory cortex (for projected target-elicited data) in the task-engaged state. For the auditory cortex, data is projected either on the sound decoding vector or the silence decoding vector. Error bars represent 95% confidence intervals. (400 cross-validations. p=<0.0025, p=<0.0025 & p=0.011;**: p<0.01,;*: p<0.05).

### Enhanced representation of target stimuli in A1 is a general feature of auditory Go/No-Go tasks

To determine whether the task-related increase in asymmetry between target and reference was a more general feature of primary auditory cortex responses during auditory discrimination, we applied our population analysis to other datasets collected during different tasks. All of these tasks used Go/No-Go paradigms (see Fig. S11a,e,i and Methods), in which the animals were presented with a random number of references followed by a target stimulus. In these different datasets, animals were required to discriminate noise bursts vs. pure tones (tone detect tasks), or categorize pure tones drawn from low, medium or high-frequency ranges (frequency range discrimination task). Contrasting datasets were obtained from two groups of ferrets that were separately trained on approach and avoidance versions of the same tone detect task. These two behavioral paradigms used exactly the same stimuli under two opposite reinforcement conditions 30, requiring nearly opposite motor responses (Fig. S11a,e). A crucial feature shared by all these tasks lies in the fact that the behavioral response to the target stimulus always required a behavioral change relative to sustained baseline activity. More specifically the target was the No-Go stimulus in negative reinforcement tasks and required animals to *cease* ongoing licking, whereas the target was the Go stimulus in the positive reinforcement task and required animals to *begin* licking in a non-lick context. In all of the analyses, lick-related neurons were removed using the approach outlined earlier.

Performing the same analyses on all tasks showed that projections of target- and reference-evoked activities in passive conditions contained a variable degree of asymmetry in the sound and silence epochs. However, in all tasks we found that task-engagement leads an enhancement of target-driven encoding during sound (Fig. 8a,b;e,f;i,j;m,n). As previously described for the rate discrimination task (Fig. 3 and 4e), target projections more strongly deviated from baseline than projections of reference stimuli in the engaged condition. Moreover, for three of the four tasks we examined, enhancement of target representations was not observed at the level of population-averaged responses, but only in the direction determined by the decoder (Fig. 8b,f,j,n). During the post-sound silence, decoding accuracy quickly decayed in both passive and engaged states, but remained above chance (Fig. S11c,g,k). As in the click-train detection task, decoding accuracy relied on a different encoding strategy than the sound period (Fig. S11d,h,l), and the asymmetry during the post-sound silence was high both in passive and engaged conditions (Fig. S12).

**Figure 8.**
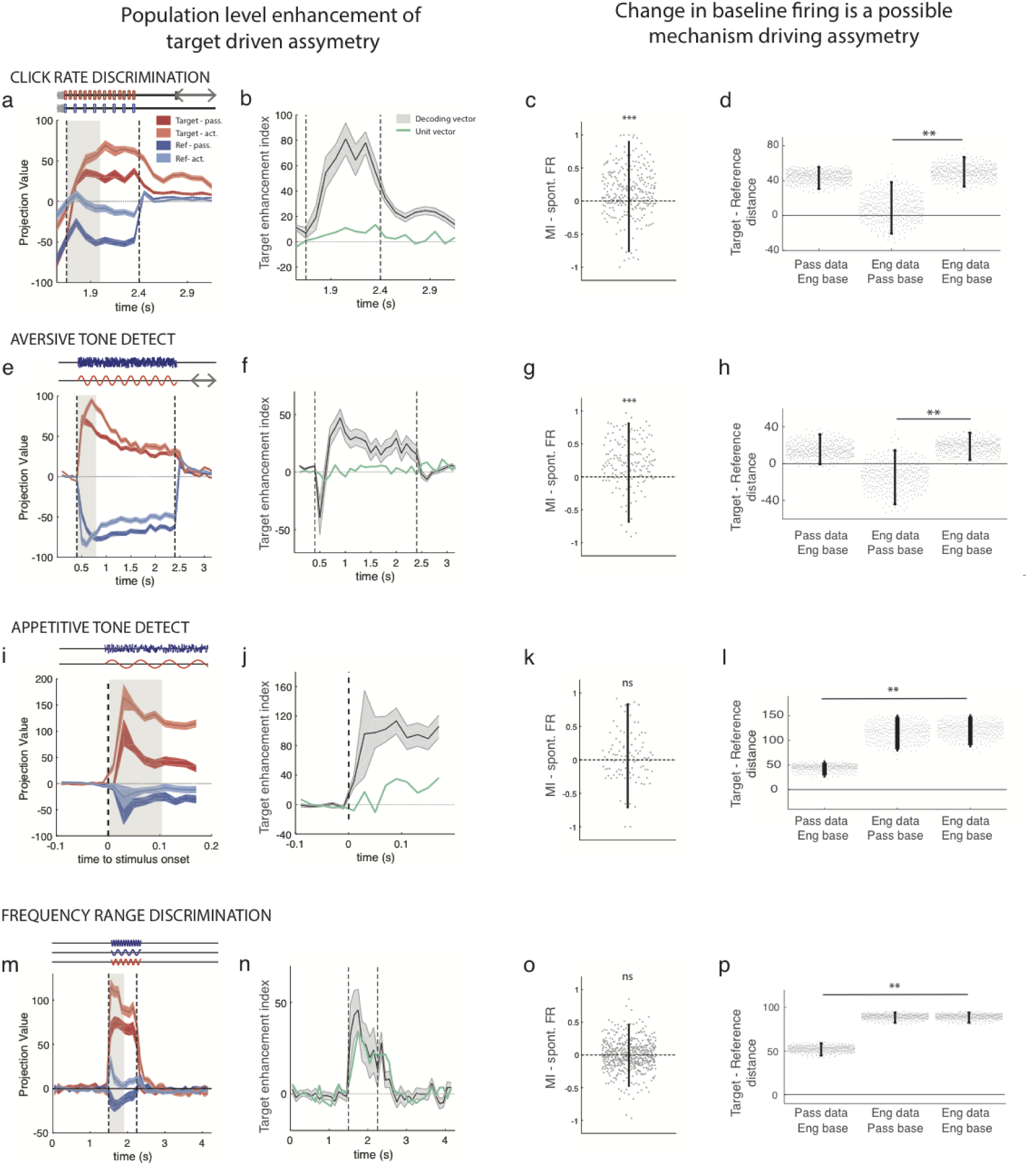
Asymmetric encoding of target and reference stimuli in a range of auditory discrimination tasks. *Each line of four panels represent the same analysis for all four tasks, statistics are given in order of appearance in the figure: click rate discrimination, aversive tone detect, appetitive tone detect, frequency range discrimination*. *a,e,l,m Projection onto the decoding axis determined during the sound period of trial-averaged reference (blue) and target (ref) activity during the passive (dark colors) and the active (light colors) sessions. A baseline value computed from pre-stimulus spontaneous activity was subtracted for each neuron, so that the origin corresponds to the projection of spontaneous activity (shown by black line). Note that the target-driven activity is further from the baseline in the active state and the reference-driven activity is closer. The periods used to construct the decoding axis are shaded in gray. Error bars represent 1 std calculated using decoding vectors from cross-validation (n=400)*. *b,f,j,n Index of target enhancement induced by task engagement based on projections using the decoding axis determined during the sound. In green same index instead giving the same weight to all units. The difference between the green and black curved indicates that the change in asymmetry induced by task engagement cannot be detected using the population averaged firing rate alone. Error bars represent 1 std calculated using decoding vectors from cross-validation (n=400)*. *c,g,k,o Modulation index of each unit for spontaneous firing rate after exclusion of lick-related units. Eror bars are 95% C.I. (one-sample two-sided Wilcoxon signed rank test with mean 0, n=277, zval=6.35, p=2.1e-10; n=161, zval=7.22, p=5.4e-13; n=99, zval=1.01, p=0.30; n=520, zval=-0.78, p=0.47; ***: p<0.001)*. *d,h,l,p Comparison of reference/target asymmetry for evoked responses in different states compared to different baselines given by passive or engaged spontaneous activity. Reference/target asymmetry is the difference of the distance of target and reference projected data to a given baseline. We examine three cases: (i) passive evoked responses, distances calculated relative to engaged spontaneous activity; (ii) engaged evoked responses, distances calculated relative to passive spontaneous activity; (iii) engaged evoked responses, distances calculated relative to engaged spontaneous activity. In all three cases, the engaged decoding axis was used for projections. Error bars represent 95% confidence intervals (n=400 cross validations; p(col1,col3)=0.29 & p(col2,col3)<0.0025; p(col1,col3)=0.38 & p(col2,col3)<0.0025; p(col1,col3)<0.0025 & p(col2,col3)=0.16; p(col1,col3)<0.0025 & p(col2,col3)=0.92; **: p<0.01)*.

Comparison of appetitive and aversive versions of the same task is particularly revealing as to which type of stimulus was associated with enhanced representation in the engaged state. In the appetitive version of the tone detect tak, ferrets needed to refrain from licking on the reference sounds (No-Go) and started licking the water spout shortly after the target onset (Go) (Fig. S11e), whereas in the aversive (conditioned avoidance) paradigm they had to stop licking after the target sound (No-Go) to avoid a shock (Fig.S11a). It is important to note that although the physical stimuli presented to the behaving animals were identical in both tone detect tasks, the associated motor behaviors of the animals are nearly opposite. Projection of task-engaged A1 population activity reveals a target-driven encoding (compare right panels of Fig. 8f,j with Fig. 8I,j), irrespective of whether the animal needed to refrain from or to start licking to the target stimulus. This shows that the common feature of stimuli that are enhanced after projection onto the decoding axis is that they are associated with a change of ongoing baseline behavior.

This range of behavioral paradigms provides additional arguments against the described changes in activity being solely due to correlates of licking activity. Firstly, we observed enhanced target-driven encoding in both the appetitive and aversive tone-detect paradigms, even though the licking profiles were diametrically opposite to each other. Secondly, comparing the projections of the population activity in the approach tone detect task with the click rate discrimination task reveals a strong similarity in the temporal pattern of asymmetry observed during task engagement. In less than 100 ms, projection of target-elicited activity reached its peak in both paradigms (Fig. 8a,i), although the direction and time course of the licking responses were reversed, with a fast decline in lick frequency for the click rate discrimination task (Fig. 1a), versus a slow increase for the tone detect (Fig. S11e left panel). Last, although the results are more variable partly due to low decoding performance, we observed target-driven encoding during the post-stimulus silence in the passive state (Fig. S12) although ferrets were *not* licking during this epoch. The points listed here are again in agreement with a representation of the stimulus’ behavioral consequences, independent of the animal motor response.

As pointed out in the case of the click rate discrimination task, the enhancement of target representation in the engaged condition can rely on two different mechanisms, a shift in the spontaneous activity or a shift in stimulus-evoked activity. We therefore set out to tease apart the respective contributions of the two mechanisms in this novel set of tasks. As in Fig. 6, we compared the distance of target and reference passive and engaged projections to either engaged or passive baseline activities. Out of the three additional datasets, we observed an increase in spontaneous firing rates only in the aversive tone detect task (Fig. 8g). In this latter paradigm, task-induced modulations of spontaneous activity patterns explained the change in asymmetry during sound presentation, similar to what was observed in the click rate discrimination task (compare Fig. 8d and 8h). The other two tasks showed no global change of spontaneous firing rate (Fig. 8k,o), and consequently, during the task engagement, the enhancement of the target representation was solely due to the second mechanism, the changes in the target-evoked responses themselves (Fig.8l,p). During the silence, we observed as previously for the click-rate discrimination that the increase in asymmetry relied only on the second mechanism (Fig. S11).

Taken all together, population analysis on four different Go/No-Go tasks revealed an increase of the encoding in favor of the target stimulus as a general consequence of task-engagement on A1 neural activity. Viewing activity changes in this light allowed us to interpret the previously observed changes in spontaneous activity as one of two possible mechanisms underlying this task-induced change of stimulus representation in A1 population activity.

## DISCUSSION

In this study, we examined population responses in the ferret primary auditory cortex during auditory Go/No-Go discrimination tasks. Comparing responses between sessions in which animals passively listened and sessions in which animals actively discriminated between stimuli, we found that task-engagement induced a shift from a sensory-driven to an asymmetric, target enhanced, representation of the stimuli, highly similar to the type of activity observed in dorsolateral frontal cortex during engagement in the same task. This enhanced representation of target stimuli was found in a variety of discrimination tasks that shared the same basic Go/No-Go structure, but used a variety of auditory stimuli and reinforcement paradigms.

In the click rate discrimination task that we analyzed first, the sustained asymmetric stimulus encoding in A1 was only observed in the engaged state (Fig. 3). One possible explanation is that this encoding scheme relied on corollary neuronal discharges related to licking activity. However there are several factors that argue against this interpretation. Firstly, we adopted a stringent criterion for the exclusion from the analysis of all units whose activity was correlated with lick events (Fig. 4). After removing lick-responsive units from the analysis the results remained unchanged, indicating the absence of a direct link between licking and the observed asymmetry in the encoding. Furthermore, the large differences in the lick profiles between the different tasks were not in line with the remarkably conserved target-driven projections of population activity across tasks and reinforcement types, supporting a non-motor nature of the stimulus encoding in A1 (Fig. 8b,f,j,n). Finally, the role of baseline shifts due to the change in spontaneous activity in two more tasks further argues against a purely motor explanation of the observed asymmetry (Fig. 6 and Fig. 8a) since the spontaneous activity occurs during epochs that preceded stimulus presentation and behavioral changes. Altogether, while the different lines of evidence exposed above make an interpretation in terms of motor activation unlikely, ultimately a different type of behavioral report, such as one using similar responses, would help fully rule out this possibility.

Our analyses show that the target-driven encoding scheme during task engagement is neither purely sensory nor purely motor, but instead argue for a more abstract, cognitive representation of the stimulus behavioral meaning in A1 during task engagement. As the target stimulus was associated with an absence of licking in the tasks under aversive conditioning, one possibility could have been that the A1 encoding scheme was contrasting the only stimulus associated with an absence of licking (No-Go) against all other stimuli (Go). This lick/no-lick encoding was however not consistent with the tone detect task under appetitive reinforcement, in which the target stimulus was a Go signal for the animal. We thus suggest that A1 encodes the behavioral meaning of the stimulus by emphasizing the stimulus requiring the animal to change its behavioral response, i.e. the target stimuli in the different tasks we examined. However, our data do not allow us to conclude whether this behavioral meaning corresponds to the encoding of the stimulus-action association, or the animal’s decision, or the output motor command leading to a change in behavioral response and it would be interesting to perform similar analyses in tasks more specifically designed to tease apart these different possible interpretations.

### Relation to previous studies

A series of previous studies found that task-engagement strongly influences responses in the primary auditory cortex, in some cases sharpening stimulus representation^26–28,37^, in others leading to a suppression of sensory responses^14^, as was also observed during locomotion^17,18^. While some studies observed signatures of decision-related activity in A1^11,38^, none has hitherto reported the strong representation of behavioral meaning described here in the population code.

The majority of previous studies concentrated on single-neuron or LFP activity. In contrast, our results critically rely on population-level analyses^39–42^, and in particular, on linear decoding of population activity. This is a simple, biologically-plausible operation that can be easily implemented by a neuron-like readout unit that performs a weighted sum of its inputs. The summed inputs to this hypothetical read-out unit showed that Go and No-Go stimuli elicited inputs symmetrically distributed around spontaneous activity in the passive state. In contrast, in the task-engaged state, only target stimuli, which required an explicit change in ongoing behavior, led to an output different from spontaneous activity, once passed through the readout unit. This switch from a more symmetric, sensory-driven to an increasingly asymmetric, target-driven representation was not clearly apparent if single-neuron responses were simply averaged or normalized (Fig. S7, 7b,f,j,n), but instead relied on a population analysis in which different units were assigned different weights by projecting population activity on the decoding axis. Note that the weights were not optimized to maximize the asymmetry between Go and No-Go stimuli, but rather the discrimination between them. The shift towards a more asymmetric representation of the behavioral meaning of stimuli is therefore an unexpected but important by-product of the analysis.

From a population-decoding viewpoint, task-engagement induced a shift towards an enhanced representation of target stimuli class in all the tasks we considered. However, considering these same effects from a less elaborate sensory coding view, they appear to be quite varied and to depend on the details of the stimuli. Thus, in the tone-detection task, previous studies reported that task-engagement enhanced the representation of the relevant tone frequency in a negative reinforcement paradigm^26–28^, and caused a suppression at the tone frequency during the appetitive version of the task^30^. In the click-discrimination task, task-engagement led to decreased temporal fidelity in the representation of click times, the main sensory features of the stimuli (see Fig. 1 and ^14^). These varied results, however, are unified by a shift to a representation of the behavioral meaning of stimuli. Our findings therefore provide a possible way to reconcile the diverse effects described earlier.

### Possible implication of an A1-FC loop during task engagement

Recordings performed in dorsolateral frontal cortex (dlFC) in the ferret during tone detection^36^ showed that, when the animal is engaged in the task, dlFC single units encode the abstract behavioral meaning of the stimuli by responding only to target stimuli (that require a change in the ongoing behavioral output) but remain silent for reference stimuli. Remarkably, projections of reference- and target-elicited A1 activity on the linear readout showed the same type of target-specific patterns of activity. Several possible mechanisms could account for these similarities of representations in A1 and dlFC. Here we propose that, during task engagement, sound evoked activity in A1 triggers activity in dlFC, which then subsequently feeds back top-down inputs to A1 that may underlie the sustained activity pattern found during post-stimulus silence.

Very early in the trial, the asymmetric encoding is already fully present in A1 (as early as 100ms in the rate discrimination task for instance; Fig. 3e top panel). At this point in time, dlFC does show some target-selective responses that increase over time (Fig. 7a). This suggests the presence of a feed-forward mechanism early in the trial, by which A1 may be feeding higher-order auditory cortex and FC with a pattern of neuronal responses encoding the behavioral meaning of the stimulus. Our results show that this early task-induced change in the representation in A1 relies on a shift of spontaneous activity at the population level that may be due to tonic top-down or neuromodulatory inputs during task engagement^43,44^. The presence of a dynamic balance characterizes interactions between A1 and dlFC has been previously shown by changes in Granger causality and effective connectivity during behavioral state transitions^45^.

As the trial progresses, the encoding in A1 progressively shifts (Fig. S6). Activity projected on the late decoding vector (Fig. 3e bottom panel) shows a progressive buildup similar to the activity observed in dlFC (Fig. 7). The late stimulus encoding, during the later phase of the click trains and the subsequent post-stimulus silence (Fig. 3e bottom panel) may thus be gradually engaging stronger top-down inputs from the dlFC-A1 network loop. The persistent encoding of stimuli identity could therefore rely on a stimulus-specific top-down input from frontal areas. Although direct connections from dlFC to A1 have not been identified in ferrets, several recent studies have identified direct inputs from the rodent motor cortex^17^, the rodent orbitofrontal cortex^46,47^ and the rodent secondary auditory areas^48^ (ferret posterior ectosylvian gyrus) to A1. Altogether, while the comparison of time-course of activity in A1 and dlFC suggest that the recruitment of the A1-FC loop is a plausible interpretation of our results, more direct evidence is needed to establish this mechanism.

### Projection to the read-out null space as a mechanism for target detection in A1

Our analysis suggests a novel population readout mechanism for extracting behaviorally relevant information from A1 while suppressing other, irrelevant sensory information: in the task-engaged state, irrelevant sensory inputs (reference stimuli) elicit changes of activity that are orthogonal to the read-out axis and therefore cannot be distinguished from spontaneous activity. This mechanism is reminiscent of the mechanism proposed for movement preparation in motor cortex^49^, where preparatory neural activity lies in the null space of the motor readout, i.e. the space orthogonal to the read-out of the motor command, and therefore does not generate movements. In our case, the readout is task-dependent, as it presumably depends on the performed discrimination task. We showed that the A1 activity in the engaged condition rearranges so that the difference between spontaneous activity and reference-elicited activity lies in the null space of the readout, which is therefore only activated by target stimuli. This rearrangement can be implemented either by a change of reference-elicited activity or by a change of spontaneous activity. In two of the examined tasks, click-discrimination and aversive tone detection, we found that the rearrangement of population activity relied mostly on the change in population spontaneous activity in the engaged condition. Strikingly, these two tasks were performed by the same ferrets, which were trained to switch between the two tasks in the same session. In the two other tasks, reference-elicited activity in the passive condition were already aligned with the passive spontaneous activity when projected on the active decoder, suggesting that learning these behavioral tasks may have profoundly reshaped stimulus-evoked activity. Our results therefore suggest that task-dependent shaping of spontaneous activity can allow the primary auditory cortex to encode the behavioral meaning of stimuli in a task-relevant, and often in a highly flexible manner.

Changes in spontaneous activity have previously been shown to contribute to stimulus responses^50–54^ and task-driven changes have been reported in multiple previous studies^14^ but, to our knowledge, have never been given a functional role in stimulus representation^55^. Here we propose that population-level modulations of spontaneous activity act as a mechanism supporting the asymmetric representation of reference and stimuli target in the engaged state. This was clearly the case in tasks where the passive reference-evoked responses and spontaneous patterns of activity were not already aligned with respect to the active decoding vector (Fig.8a-d and Fig8e-h). In those tasks, significant adjustments in spontaneous activity supported the deployment of a reference/spontaneous space orthogonal to the active readout-out axis.

However, this proposed simple linear readout mechanism cannot fully account for the whole set of responses observed in frontal areas for at least two reasons. First, projections of reference-elicited activity (in A1) during engagement on an aversive task still give rise to a non-null, albeit reduced, output contrary to what is observed in dlFC area recordings. Second, projecting passive data onto the engaged decoding vector results in symmetric and reduced outputs (data not shown), whereas dlFC recordings showed on average a complete absence of response during passive state during the tone-detect task^36^. An additional non-linear gating mechanism likely operates between primary auditory cortex and frontal areas, further reducing responses to any stimulus in the passive state and to reference sounds in the active state. In particular, neurons in higher-order auditory areas could refine the population-wide, abstracted representation originating in A1 through the proper combinations of synaptic weights. Such a mechanism could also explain why individual single units recorded in belt areas of the ferret auditory cortex show a gradual increase in their selectivity to target stimuli^56^.

### Effects of learning

All the recordings analyzed here were performed on highly trained animals. Several investigations have reported that training procedures strongly influence neural representations in primary cortices^57–61^. One may therefore wonder to what extent our findings, even in the passive state, depend on the prior training history of the animal^62–64^. To address this question, we examined A1 recordings performed in a naive ferret exposed to the same stimuli as used in the click-train discrimination task. Stimulus discrimination was relatively decreased, during both the sound and silent periods when compared with the decoder accuracy obtained with trained animals (Fig. S5c,d). In particular, the discrimination performance during the post-stimulus silence was reduced to chance-levels, while in trained animals it was above chance even in the passive state. The weak but significant maintained encoding of stimulus class observed in the passive state with expert ferrets thus appears to be due to the behavioral training. Discrimination in the passive condition for trained animals also involved target-specific activity during post-stimulus silence (Fig. 3c,d, bottom panels), whereas it was not the case for naive ferrets (Fig. S5d), indicating that this target-driven mechanism is ubiquitously present during the silent period in trained animals.

Interestingly, passive projections of target- and reference-evoked activities showed variable degrees of asymmetry across tasks (Fig. 3c and 8a,e,i,m). This observation could be explained by the variability in training duration across ferrets, in task performance, and in paradigm requirements and complexity. Strikingly, the only task we examined involving long-term memory (frequency range discrimination task) exhibited a very strong asymmetry *both* in passive and active states (Fig. 8m). While asymmetric representation of stimuli was weak in tasks demanding flexible and rapid attention towards new stimuli (rate discrimination and tone detect tasks), a task involving long-term memory, such as the frequency range discrimination task, could engage global reshaping of the neuronal population structure to keep a mnemonic trace of the behaviorally-relevant stimuli. Interestingly, this target-driven asymmetry in the passive state came along with a lack of change in the spontaneous population activity between passive and active state (Fig. 8fo). This observation is in agreement with the hypothesis that the encoding of stimulus behavioral meaning is mediated by an adjustment of spontaneous population activity, mostly operated in passive state for this particular task.

In summary, we found that task-engagement induces a shift from sensory-driven to abstract, behavior-driven representations in the primary auditory cortex. These abstract representations are encoded at a population, but not at a single-neuron level, and strikingly resemble abstract representations observed in higher-level cortices. These results suggest that the role of primary sensory cortices is not limited to encoding sensory features. Instead, primary cortices appear to play an active role in the task-driven transformation of stimuli into their behavioral meaning and the translation of that meaning into task-appropriate motor actions.

### Materials and methods

### Training and recordings

#### Behavioral training

All experimental procedures conformed to standards specified by the National Institutes of Health and the University of Maryland Institutional Animal Care and Use Committee (IACUC). Adult female ferrets, housed in pairs in normal light cycle vivarium, were trained during the light period on a variety of different behavioral paradigms in a freely moving training arena. After headpost implantation, the ferrets were retrained while restrained in a head-fixed holder until they reached performance criterion again. Most of the animals in these studies were trained on multiple tasks, including the two ferrets trained both on the click rate discrimination and the tone detect tasks. Three out of four tasks shared the same basic structure of Go/No-Go avoidance paradigms^65^, in which ferrets were trained in a conditioned avoidance paradigm to lick water from a spout during the presentation of a class of reference stimuli and to cease licking after the presentation of a different class of target stimuli to avoid a mild shock. The positive reinforcement task is detailed below (see *Tone detect task* – *Aversive conditioning*).

Recordings began once the animals had relearned the task in the holder. Each recording session included epochs of passive sounds presentation without any behavioral response or reinforcement, followed by an active behavioral epoch where the animals could lick. A post-passive epoch was then recorded. This sequence of epochs could be repeated multiple times during a recording session. The table below summarizes the animals and recordings for each task.

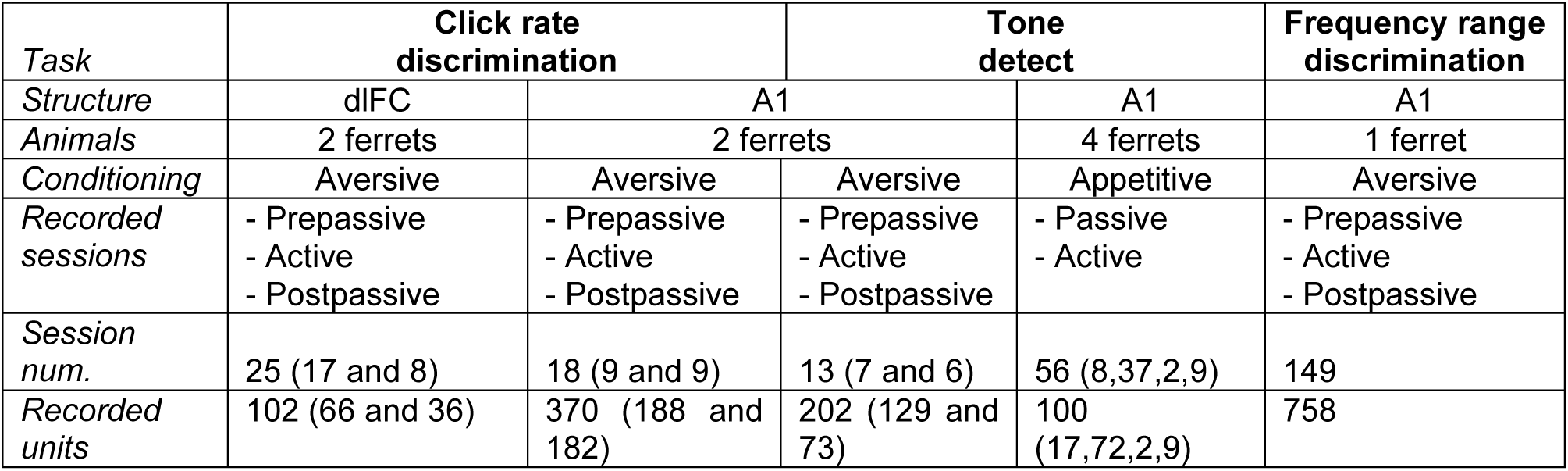

#### Click rate discrimination task

Two adult female ferrets were trained to discriminate low from high rate click trains in a Go/No-Go avoidance task. A block of trials consisted of a sequence of a random number of reference click train trials followed by a target click train trial (except on catch blocks in which 7 reference stimuli were presented with no target). On each trial, the click train was preceded by a 1.25s neutral noise stimulus (Fig. 1A). Ferrets licked water from a spout throughout trials containing reference click trains until they heard the target sound. They learned to stop licking the spout either during the stimulus or after the target click train ended, in the following 0.4-s time silent response window, in order to avoid a mild shock to the tongue in a subsequent 0.4 s shock window (Fig.1A). Any lick during this shock window was punished. The ferrets were first trained while freely-moving daily in a sound-attenuated test box. Animals were implanted with a headpost when they reached criterion, defined with a Discrimination Ratio (DR)>= 0.64 where DR = HR * (1-FA) [Hit Rate, HR=0.8 and False Alarm, FA=0.2]. They were then retrained head-fixed with the shocks delivered to the tail. The decision rule was reversed in the 2 animals, as low rates were Go stimuli for one animal and No-Go for the second one. During each session, rates were kept identical, but were changed from day to day.

#### Tone detect task – Aversive conditioning

The same two ferrets were trained on a tone detect task previously described^26^. Briefly, a trial consisted of a sequence of 1 to 6 reference white noise bursts followed by a tonal target (except on catch trials in which 7 reference stimuli were presented with no target). The frequency of the target pure tone was changed every day. The animals learned not to lick the spout in a 0.4 s response window starting 0.4 s after the end of the target. The ferrets were trained until they reached criterion, defined as consistent performance on the detection task for any tonal target for two sessions with >80% hit rate accuracy and >80% safe rate for a discrimination rate of >0.65.

#### Tone detect task – Appetitive conditioning

4 ferrets were on an appetitive version of the tone detect task previously described^30^. On each trial, the number of references presented before the target varied randomly from one to four. Animals were rewarded with water for licking a water spout in a response window 0.1–1.0 s after target onset. False alarms were punished with a timeout when ferrets licked earlier in the trial before the target window. The average DR during experiments was 0.76. This data set contained sessions with different trial durations, therefore we analysed separately data from the first 200ms after stimulus onset and 200ms before stimulus offset. For this task, the passive data was not structured in the format of successive reference and target trials as in the engaged session but instead the animal was presented with a block of reference only trials followed by a block of target only trials separately. This slight change in the structure of the sound presentation did not affect our results that were highly similar to other tasks but may explain the slightly higher accuracy of decoding during the initial silence in the passive data. Indeed reference and target trials were systematically preceded by other reference and target trials, possibly allowing the decoder to discriminate using remnant activity from the previous trial.

#### Frequency range discrimination task

One ferret was trained on a three-frequency-zone discrimination task with a Go/No-Go paradigm. The three frequency zones were defined once and for all and the animal had to learn the corresponding frequency boundaries (Low-Medium: ~500 Hz / Medium-High: ~3400 Hz). Each trial consisted of the presentation of a single pure tone (0.75-s duration) with a frequency in one of the three zones. A trial began when the water pump was turned on and the animal licked a spout for water. The ferret learned to stop licking when it heard a tone falling in the Middle frequency range in order to avoid punishment (mild shock) but to continue licking if the tone frequency fell in either the Low or High range. The shock window started ms after tone offset and lasted ms. The pump was turned off 2 s after the end of the shock window. The learning criterion was defined as DR>40% in three consecutive sessions of more than trials.

#### Acoustic stimuli

All sounds were synthesized using a 44 kHz sampling rate, and presented through a free-field speaker that was equalized to achieve a flat gain. Behavior and stimulus presentation were controlled by custom software written in Matlab (MathWorks).

#### Click rate discrimination task

Target and reference stimuli were preceded by an initial silence lasting 0.4 s followed by a 1.25 s-long broadband-modulated noise bursts (temporal orthogonal ripple combinations, TORC^66^) acting as a neutral stimulus, without any behavioral meaning (Fig.1A). Click trains all had the same duration (0.75 s, 0.8 s inter-stimulus interval of which the last 0.4 s consisted of the response window) and sound level (70 dB SPL). Rates used were comprised between 6 and 36 Hz (ferret A: references [6 7 8 15] Hz, targets [24 26 28 30 32 33 36] Hz / ferret L: references [26 28 30 32 36] Hz, targets [6 8 9 16] Hz).

#### Tone detect task

Reference sounds were TORC instances. Targets were comprised of pure tone with frequencies ranging from 125–8000 Hz. Target and reference stimuli were preceded by an initial silence lasting 0.4 s. Target and reference stimuli all had the same duration (2 s, 0.8 s inter-stimulus interval whose last 0.4 s consisted of the response window for the aversive tone detect task) and sound level (70 dB SPL). In the appetitive version of this paradigm, target and reference duration varied between sessions (0.5–1.0 s, 0.4–0.5-s interstimulus interval).

#### Frequency range discrimination task

The target frequency region was the Medium range (tone frequencies: 686, 1303 and 2476 Hz) while the reference regions were the Low and High frequency ranges (100, 190 and 361 Hz; 4705, 8939 and 16884 Hz). Thus the set of tones included 9 frequencies with 90% increment (~0.9 octave) and spanned a ~7.4 octaves range. Target and reference stimuli (duration: 0.75 s; level: 70 dB SPL) were preceded by an initial silence lasting 1.5 s and followed by a 2.4 s silence comprising the shock window (400 ms starting 100 ms after the tone offset).

#### Neurophysiological recordings

To secure stability for electrophysiological recording, a stainless steel headpost was surgically implanted on the skull (Fritz et al. 2003; Fritz et al. 2010). Experiments were conducted in a double-walled sound attenuation chamber. Small craniotomies (1–2 mm diameter) were made over primary auditory cortex prior to recording sessions, each of which lasted 6–8 h. The A1 and frontal cortex (dorsolateral FC and rostral ASG) regions were initially located with approximate stereotaxic coordinates and then further identified physiologically. Recordings were verified as being in A1 according to the presence of characteristic physiological features (short latency, localized tuning) and to the position of the neural recording relative to the cortical tonotopic map in A1^67^. Data acquisition was controlled using the MATLAB software MANTA^68^. Neural activity was recorded using a 24 channel Plexon U-Probe (electrode impedance: ~ 275 kΩ 1 kHz 75-μm inter-electrode spacing) during the click discrimination task and the aversive version of the tone detect task. Recordings during the other tasks (frequency range discrimination and appetitive tone detect task) were done with high-impedance (2-10 MΩ) tungsten electrodes (Alpha-Omega and FHC), using multiple independently moveable electrode drives (Alpha-Omega) to independently direct up to four electrodes. The electrodes were configured in a square pattern with ~800 μm between electrodes. The probes and the electrodes were inserted through the dura, orthogonal to the brain’s surface, untile the majority of channels displayed spontaneous spiking.

### Data Analysis

Data analyses were performed in MATLAB (Mathworks, Natick, MA, USA).

#### Spike sorting

To measure single-unit spiking activity, we digitized and bandpass filtered the continuous electrophysiological signal between 300 and 6,000 Hz. The tail shock for incorrect responses introduced a strong electrical artefact and signals recorded during this period were discarded before processing.

Recordings performed with 24 channel Plextrodes (U-probes) (click discrimination and the tone detect tasks) were spike sorted using an automatic clustering algorithm (KlustaKwik,^69^), followed by a manual adjustment of the clusters. Clustering quality was assessed with the isolation distance, a metrics developed by Harris et al, 2001 which quantifies the increase in cluster size needed for doubling the number of samples. All clusters showing isolation distance larger than 20 were considered as single units^70,71^. A total of 82 single units and 288 multi-units were isolated. All analyses were reproduced on both pools of units and qualitatively similar results were obtained (Supplementary Information). We thus combined all clusters for the analysis. Spike sorting was performed on merged data sets from pre-passive, active and post-passive sessions.

For recordings performed with high-impedance tungsten electrodes (frequency range discrimination and relative pitch tasks), single units were classified using principal components analysis and k-means clustering followed by manual adjustment^26^.

#### Depth determination in the click rate discrimination task

Each penetration of the linear electrode array produced a laminar profile of auditory responses in A1 across a 1.8 mm depth. Supra- and infragranular layers were determined with LFP responses to 100 ms tones recorded during the passive condition. The border between superficial and middle-deep layer was defined as the inversion point in correlation coefficients between the electrode displaying the shortest response latency and all the other electrodes in the same penetration^72,73^.

#### Click reconstruction from neural data

Optimal prior reconstruction method^74^ was used to reconstruct stimulus waveform from click-elicited neural activity. Units with spontaneous firing rate larger than 2 spikes/s in at least one condition were considered for this analysis. Neuronal activity was binned at 10 ms in time with a 1-ms time step. For each trial, we defined *S*^k^(*t*) S^k^(t) the stimulus waveform of trial *k* (t ∈ [1,T]) and 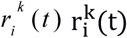 the binned firing rate of each neuron *i* ∈ [1,N] where t ∈ [1,T+τ] with τ the considered delay in the neuronal response. A linear mapping was assumed between the neuronal responses and the stimulus:

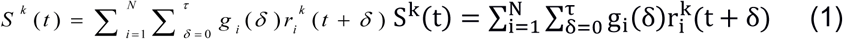

for unknown coefficients *g_i_*(*δ*). Equation (1) was rewritten as:

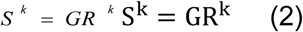

with 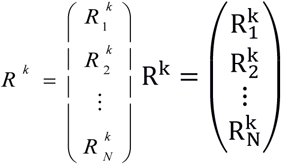 and 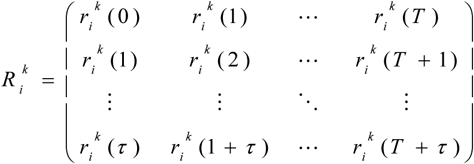 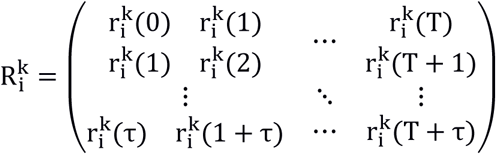 the lagged neuronal response, *G* = (*G*_1_, *G*_2_ … *G_N_*) G = (G_1_, G_2_ … G_N_) and *G_i_* = (*g_i_*(0), *g_i_*(1) … *g_i_*(*τ*)) G_i_ = (g_i_(0) g_i_(1) … g_i_(τ)) the corresponding reconstruction filter. The estimate 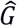 is produced by least-square fitting

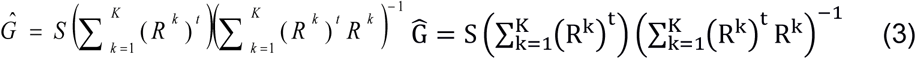

Before the inversion in the previous formula, a single value decomposition was used to eliminate the noisy components of the auto-correlation matrix. The maximal number of components retained was empirically set to 70. Once the values 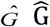 were fitted on all the trials but one, the reconstructed stimulus 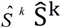 was defined as 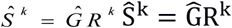 with the neuronal response *R* of the remaining run. Each trial was left out in turn. Reconstruction error was quantified with the mean-squared error (MSE) of the reconstructed stimulus. One passive and active reconstruction filters were fitted for each type of stimulus (reference and target) in every session.

#### Modulation index

To evaluate changes in a given parameter X (firing rate, vector strength) at the level of the individual unit, we define the modulation index to compare situation 1 and 2 as for each neuron as:

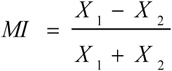

As a measure of the enhancement of target projection relative to reference projection in the task engaged state we used the following index (referred to target enhancement index in the text)

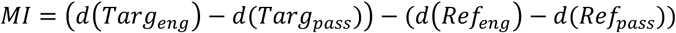

where d is the distance from baseline.

When simply measuring the asymmetry between reference and target in condition X, we used the following index (Fig. 5b; 7d,h,l,p; S9c,f,i):

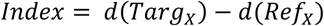

#### Vector strength

Vector strength (VS) allows to measure how tightly spiking activity is locked to one phase of a stimulus. If all spikes at exactly the same phase, VS is one whereas if firing is uniformly distributed over phases VS is 0. It is defined in Goldberg & Brown 1969 as

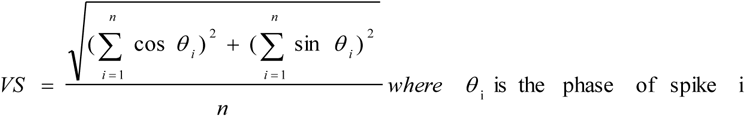

Significance was assessed using Rayleigh’s statistic, p= e^nr2^, where r is the vector strength and used p < 0.001 as the criterion for significant phase locking consistent with previous work^76^.

#### Linear discriminant classifier performance

To evaluate the accuracy with which single-trial population responses could be classified according to the presented stimulus (reference or target), we trained and tested a linear discriminant classifier^39,77^ using cross validation (FigS3).

Trial by trial pseudo-population firing rate vectors were constructed for each 100ms time bin using units from all sessions and both animals. Training and testing sets were constructed by randomly selecting equal numbers (15) of reference and target trials for each unit. All contribution of noise correlations among neurons are therefore destroyed by this procedure as the pseudo-population vector contains activity of units recorded on different days and on different trials. Since correlations between neurons can affect population coding ^33^ and are modified by task engagement ^13^,

The classifier was trained for each time bin using the average pseudo-population vectors *c*_R,t_ and *c*_T,t_ calculated from a random selection of an equal number of reference and target trials. These vectors define at time bin t the decoding vector w_t_ given by

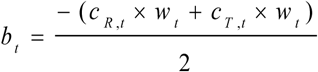

and the bias bt given by

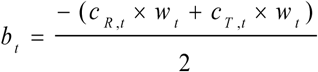

we also used Fisher discriminant analysis in which the decoding vector is defined as:

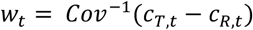

where Cov is the covariance matrix, which allows to correct the decoding vector by taking into account the trial by trial correlations between units

These define the decision rule for a new population vector x,

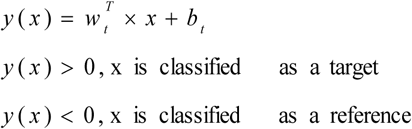

This rule was applied to an equal number of reference and target testing trials drawn from the remaining trials that were not used to train the classifier. The proportion of correctly classified trials gave the accuracy of the classifier. Cross-validation was performed 400 times by randomly picking training and testing data to estimate the average and variance of accuracy. This allowed comparing the performance of classification in two behavioral states by constructing confidence intervals from the cross-validation. Note that this limits p-value estimate to a minimum of 1/400=0.0025.

#### Random performance

To evaluate whether the classifier performance is higher than chance, the classifier was trained and tested on surrogate data sets constructed by shuffling the labels (‘reference’ and ‘target’) of trials. For each of 100 label permutations, cross-validation was performed 100 times. This allows comparing the performance of classification with chance levels by constructing confidence intervals from the cross-validation and from the random shuffled permutations.

#### Classifier evolution

When studying the evolution of population encoding (Fig. S6), we defined early sound, late sound, and silence periods as 1700-1900 ms, 2200-2400 ms and 2700-2900 ms (equal duration for comparison) relative to trial onset. The classifier was trained on randomly chosen trials from one time period and then tested on trials at all other 100ms time bins.

We also constructed matrices showing the accuracy of the classifier trained and tested at all 100ms time bins and evaluated whether these values are higher than chance using surrogate data sets by shuffling labels as described above.

When comparing the classifier during sound and silence periods across tasks (Fig. 7), the following periods were used:

|  | Click rate discrimination | Aversive tone detect | Appetitive tone detect | Frequency range discrimination |
| --- | --- | --- | --- | --- |
| Sound | 1.6-2s | 0.4-0.8s | 0-0.1 after stim onset | 1.5-1.9s |
| Silence | 2.5-2.9s | 2.5-2.9s | 0-0.1 after stim offset | 2.4-2.8s |

#### Projection onto decoding vectors

To study the contribution of reference and target trials to classifier performance, we projected population firing vectors at each time bin onto decoding vectors calculated during the sound and silence periods as defined above. Before projection, the mean spontaneous activity of each unit was subtracted from its firing rate throughout the whole trial. Deviations from 0 of the projection show activity deviating from spontaneous activity along the decoding axis.

#### Controlling for lick-responsive neurons

In order to control for the contribution of units directly linked with task-related motor activity to our results, we combined reconstruction and decoding methods to identify and remove lick-responsive neurons so that linear classification no longer yielded any licking-related information. The approach comprised the following steps:

- Optimal prior reconstruction (described in *Click reconstruction from neural data*) was used to reconstruct lick-activity separately for each unit.
- Reconstruction values for each unit were then sampled at the time of licks and at randomly selected times without licking. These values were used to construct population vectors of lick and non-lick activity.
- A linear classifier (described in *Linear discriminant classifier performance*) was trained and tested using cross-validation to distinguish lick from non-lick events.
- Reconstruction values and classification was also performed on random data obtained by reconstructing the licking activity of a session with the neural activity of a subsequent session. This made it possible to establish the distribution of accuracy for randomized data.
- The accuracy of classification was compared between the true data and the randomized data sets and a p-value was calculated by counting the number of permutations showing better accuracy for the randomized data than the true data.
- We progressively removed units, starting with those with highest classifier weights, which reduced the accuracy of classification, until the p-value of population classification rose above 0.4. This indicated that the remaining units contained no more information about lick events than randomized data.
- Only the units remaining after this procedure were used to re-analyze the data and verify that reliable classification and difference in projections of reference and tone trials did not rely on the difference in licking activity between the two trials.
For the click rate discrimination task only a subset of sessions (15/18) had reliable recordings of all lick events, so the analysis was done on 308 units (not 370), 277 units were identified as non-lick related. For the appetitive tone task 99/100 units, for the aversive tone task 161/202 and for the frequency range discrimination 520/758.

#### Gaussian-process factor analysis

To visualize neural trajectories of the large population of units recorded in A1, we used Gaussian-process factor analysis as described in ^78^. This method has the advantage over more traditional methods of dimensionality reduction such as PCA of jointly performing both the binning/smoothing steps and the dimensionality reduction.

#### Statistics

Statistics on classifier performance relied on p-value estimation using cross-validation. For each statistical analysis provided in the manuscript, the Kolmogorov–Smirnov normality test was first performed on the data. As the data failed to meet the normality criterion, statistics relied on non-parametric tests. When performing systematic multiple tests, the Bonferroni correction was applied.

#### Data availability

The data that support the findings of this study are available from the corresponding author upon reasonable request.

#### Code availability

Code used in the article can be supplied upon request by writing to the corresponding author.

## Supplementary information

### Comparison of results in single and multiunits

All analyses in the main section of the paper concerning the click train discrimination task combine results from single units (isolation distance > 20, see Methods) and multi-units because we found no differences concerning their general properties (see table 1) and the main population-level results of the paper (see table 2) were maintained using SU activity only, although the power of the analysis was of course reduced.

**Table 1.** Comparison of unit properties for single and multi units. Mean +/− s.e.m are given for each value and the comparison between SU and MU is performed using a ttest. For modulation indexes (first three lines), the significance compared to zero is given in brackets. These results are identical to those found in the main paper.

|  | SU | MU | Comparison |
| --- | --- | --- | --- |
| MI : baseline | 0.14 +/- 0.03 (***) | 0.19 +/- 0.02 (***) | p=0.22 |
| MI : evoked | 0.04 +/- 0.05 (ns) | - 0.05 +/- 0.06 (ns) | p=0.22 |
| MI : vector strength | 0.05 +/- 0.006 (***) | 0.04 +/- 0.0075 (***) | p=0.25 |
| Ref FR pass. – Snd | 7.45 +/- 0.70 | 6.67 +/- 0.88 | p=0.48 |
| Ref FR eng. – Snd | 9.14 +/-0.86 | 8.38 +/- 1.06 | p=0.57 |
| Targ FR pass. – Snd | 7.78 +/- 0.72 | 6.15 +/- 0.77 | p=0.12 |
| Tar FR eng. – Snd | 9.9 +/- 0.93 | 7.9 +/- 0.97 | p=0.15 |
| Ref FR pass. – Sil | 6.34 +/-0.64 | 5.3 +/- 0.68 | p=0.25 |
| Ref FR eng. – Sil | 7.96 +/-0.76 | 7.65 +/- 0.94 | p=0.79 |
| Targ FR pass. – Sil | 6.31 +/-0.67 | 5.4 +/- 0.76 | p=0.36 |
| Targ FR eng – Sil. | 8.56 +/-0.84 | 7.26 +/- 0.99 | p=0.32 |

To verify that the population-level results were maintained SU data, despite the reduced number of units (82 SU units, 370 total units used in main paper), we recapitulate below the main results using SU activity alone.

**Table 2.** Recapitulation of important results using SU activity alone.

|  | Mean [C.I.] – signif. of comparison |
| --- | --- |
| Sound accuracy pass. and eng. | 0.97 [0.95:0.99] - 0.98 [0.94:1] **NS** |
| Silence accuracy pass. and eng. | 0.59 [0.52:0.66] - 0.78 [0.69:0.87] * |
| Sound: ref and target projected values pass. | 29 [25:36] - 26 [18:33] **NS** |
| Silence: ref and target projected values pass. | 6 [4:8] - 12 [6:16] **NS** |
| Sound: ref and target projected values eng. | 16 [8:23] - 44 [33:55] ** |
| Silence: ref and target projected values eng. | 2 [0.7:4] - 37 [33:42] ** |

We found that the significant increase in accuracy during the silence with task engagement was maintained after restriction to SU activity. We also observed the significantly greater role played by target evoked activity in the engaged state after projection (as in Fig. 3) using SU activity alone (p<0.0025). The only difference with results given in the main paper is that in the passive state during the silence the stronger contribution of target activity did not achieve significance as in Fig. 3d, bottom.

### Population-encoding dynamics change between conditions

In the analyses reported in the main text, we trained a classifier at each time point in the trial, and used it to evaluate stimulus discrimination at the same time point in held-out trials. To assess how much the underlying encoding changes over the trial, we used two procedures. First, we directly compared the classifiers determined at different time-bins by computing the correlation between them (Fig. S6a,c). Second, we used the classifier obtained at three different trial epochs (early and late stimulus, post-stimulus silence) to classify the neural activity along the whole trials (Fig. S6b,d). If the encoding of stimulus underlying stimulus discrimination changes over time in the trial, a classifier trained on one time point will lead to a lower discrimination performance at other times.

In the passive condition, we found that changes in encoding over time are weak. The encoding was highly homogeneous within stimulus presentation and during the post-sound silence (Fig. S6a). Consistent with this view, classifiers trained during the early or the late phases of the stimulus presentation could be used efficiently at all other times during stimulus presentation without an appreciable drop in accuracy (Fig. S6b, brown and orange curves). In contrast, the same classifier led to chance-level discrimination at time points after stimulus presentation. Conversely a classifier trained after stimulus presentation led to chance-level performance during stimulus presentation (Fig. S6b, yellow curve). In the passive condition, the neural encoding that underlies stimulus discrimination therefore appears to change very little during stimulus presentation, and shifts abruptly afterwards.

A different picture emerged when animals were engaged in the task. The encoding appeared to change more progressively over the trial (Fig. S6c), and a classifier trained at one point systematically led to reduced discrimination performance at other time points (Fig. S6d). Moreover, no sharp transition was apparent at the time the stimulus was switched off. In particular, a classifier trained during the stimulus presentation led to a significant discrimination performance after stimulus presentation (Fig. S6d, brown and orange curves). Conversely, a classified determined during the post-sound silence led to an above chance and progressively increasing discrimination performance during stimulus presentation (Fig. S6d, yellow curve).

Altogether, in the engaged condition, the population encoding underlying stimulus discrimination therefore appeared to progressively shift from a representation purely along a stimulus-driven axis, where categorical information was present but uncorrelated with behavior (Fig. 3c top panel), to a representation along a decision-related axis, which was directly correlated with the behavioral action (Fig. 3e bottom panel).

## Supplementary Figures

**Fig S1.**
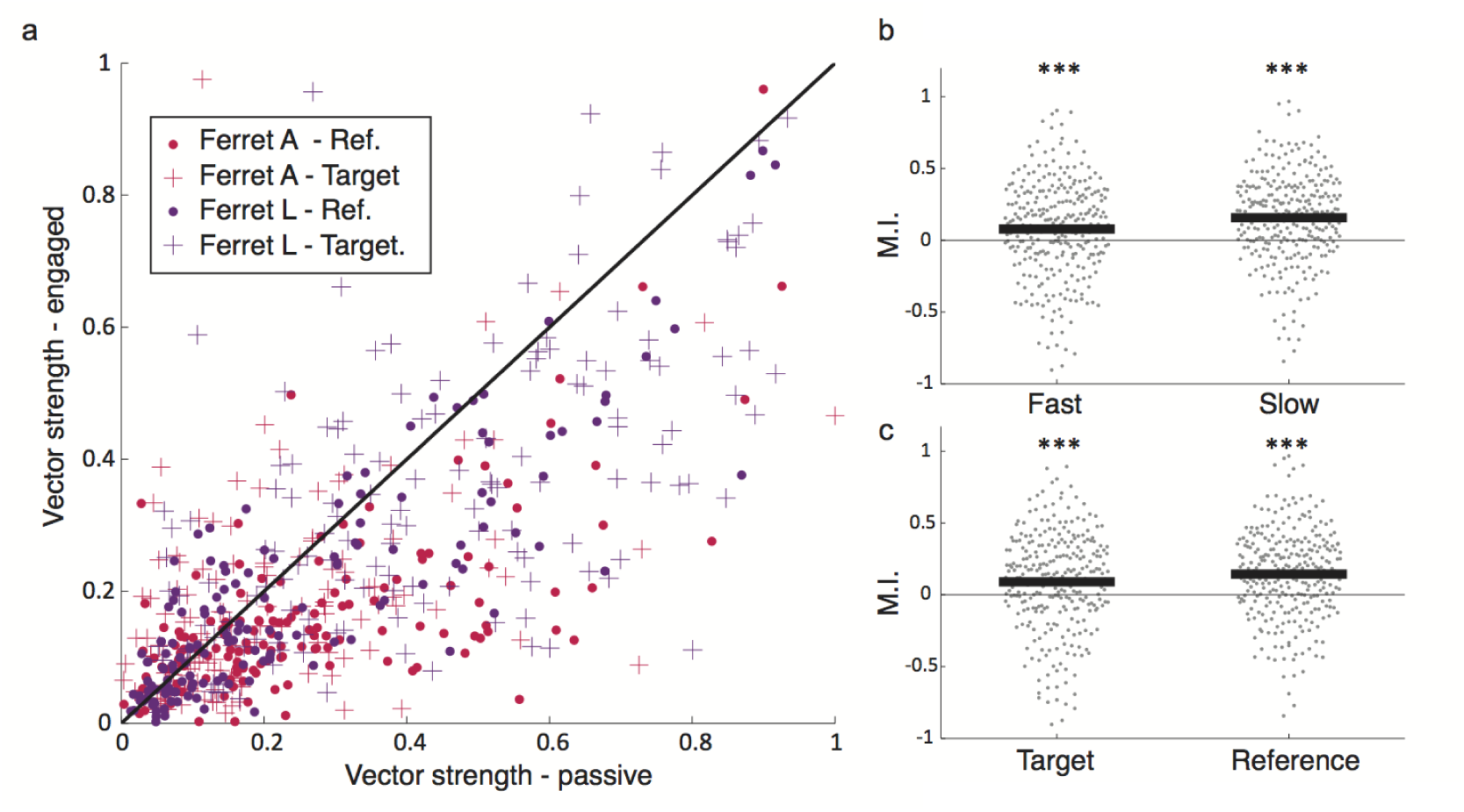
Changes in stimulus entrainment between passive and engaged conditions. *a. For each unit the vector strength for the reference and target click train is plotted in the engaged state vs the passive state. Animals are given in different colours and stimuli as different markers. Note that most points are below the x=y line, showing higher phase locking in the passive state*. *b. Modulation index of vector strength in task-engaged and passive states for fast and slow stimuli separately. (one-sample two-tailed Wilcoxon signed rank with mean 0, n=287; zval=-4.29, p=1.75e-5 & zval=-8.20, p=2.36e-16; ***: p<0.001)*. *c. Modulation index of vector strength in task-engaged and passive states for reference and target stimuli separately. (one-sample two-tailed Wilcoxon signed rank with mean 0, n=287; zval=-4.95, p=7.37e-7 & zval=-7.54, p=4.75e-14;***: p<0.001)*.

**Fig S2.**
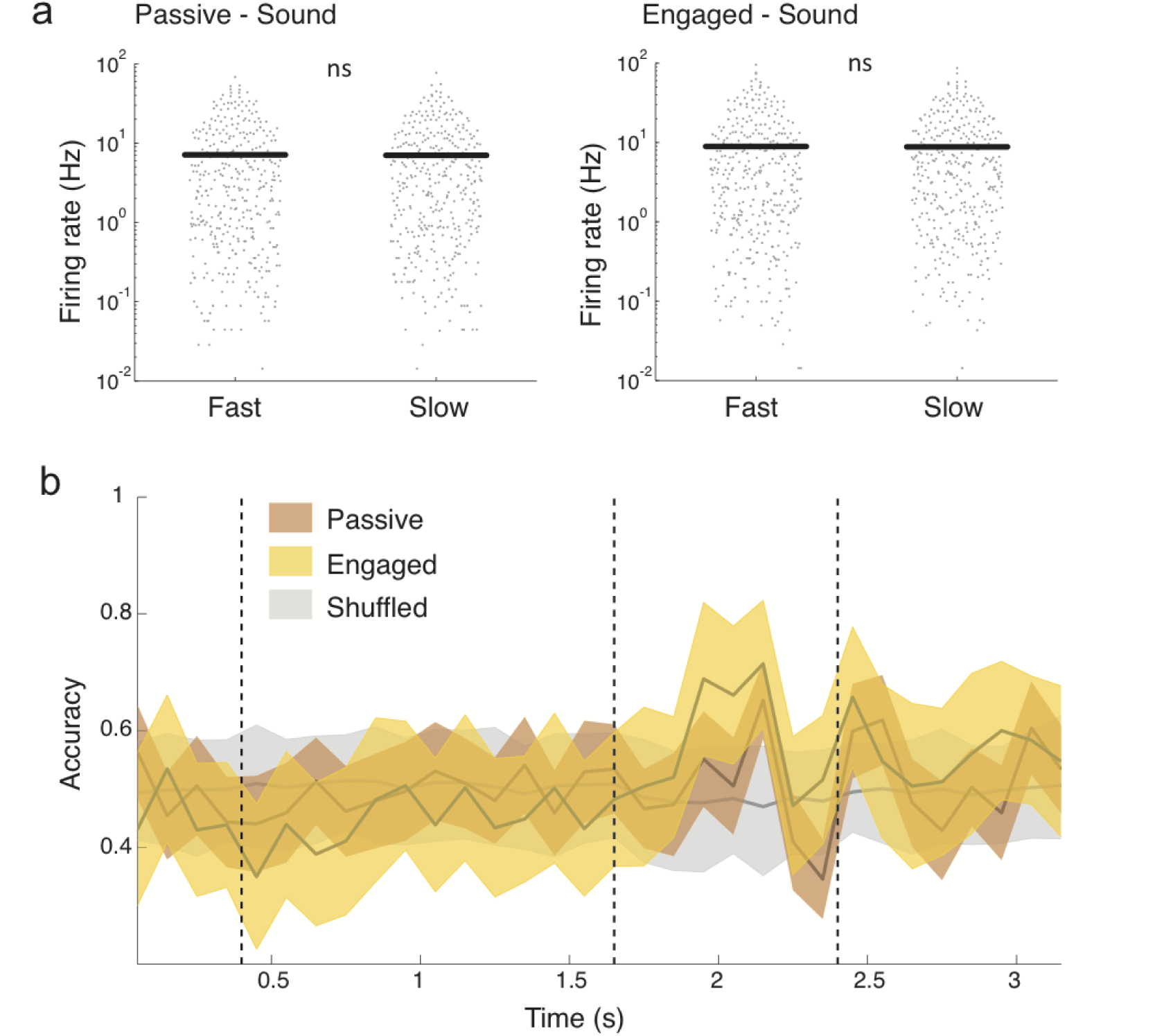
Reference and target stimuli cannot be discriminated on the basis of population-averaged activity. *a. Comparison of average firing rates on log scale in passive (left) and engaged (right) between fast and slow stimuli during the sound. (one-sample two-tailed Wilcoxon signed rank with mean 0, n=360; zval=-0.53, p=0.59 & zval=-0.25, p=0.8)*. *b. Accuracy of decoding in engaged and passive state using equal weights for all units. In grey, chance level performance evaluated on label-shuffled trials. Error bars are 1 std over 400 cross-validations*

**Fig S3.**
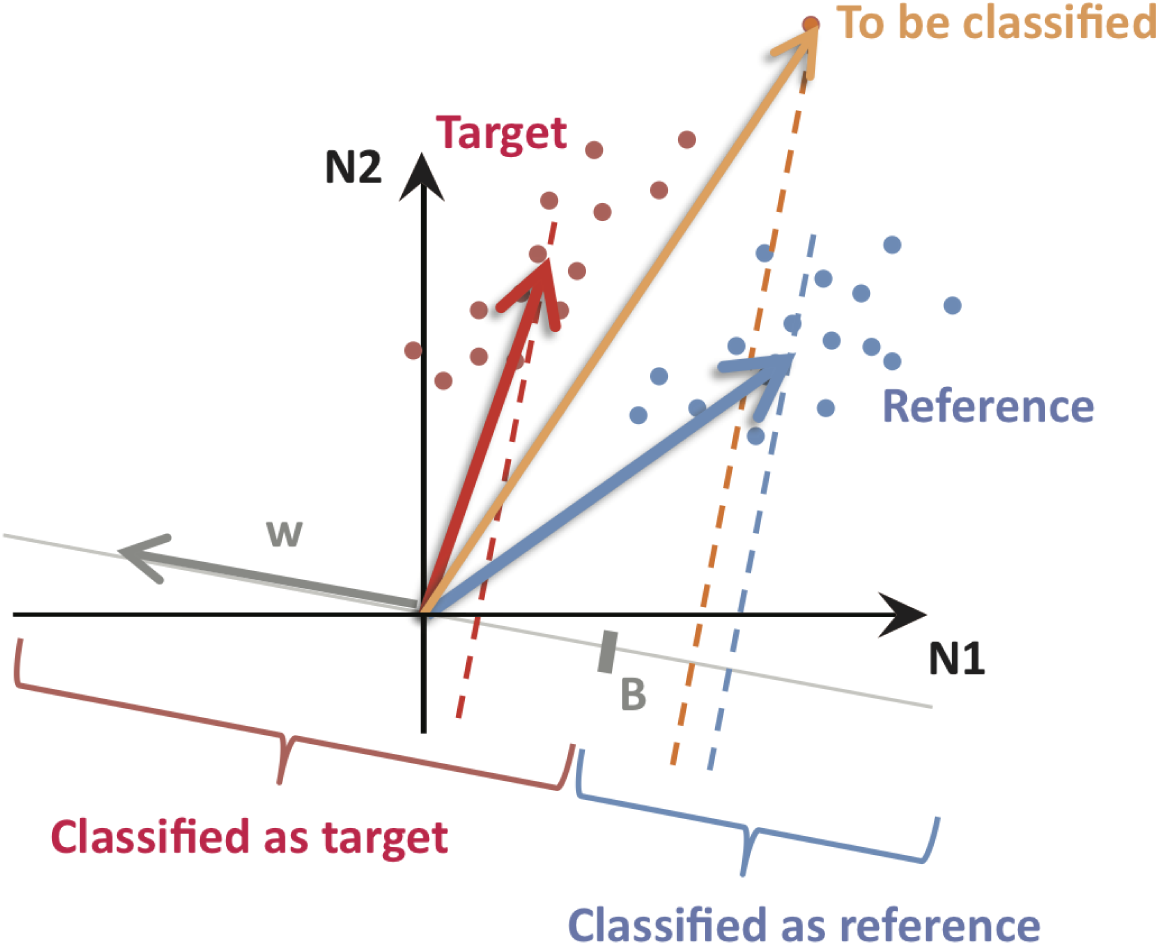
Illustration of binary classifier. *Illustration of binary classifier, see materials and methods*.

**Fig S4.**
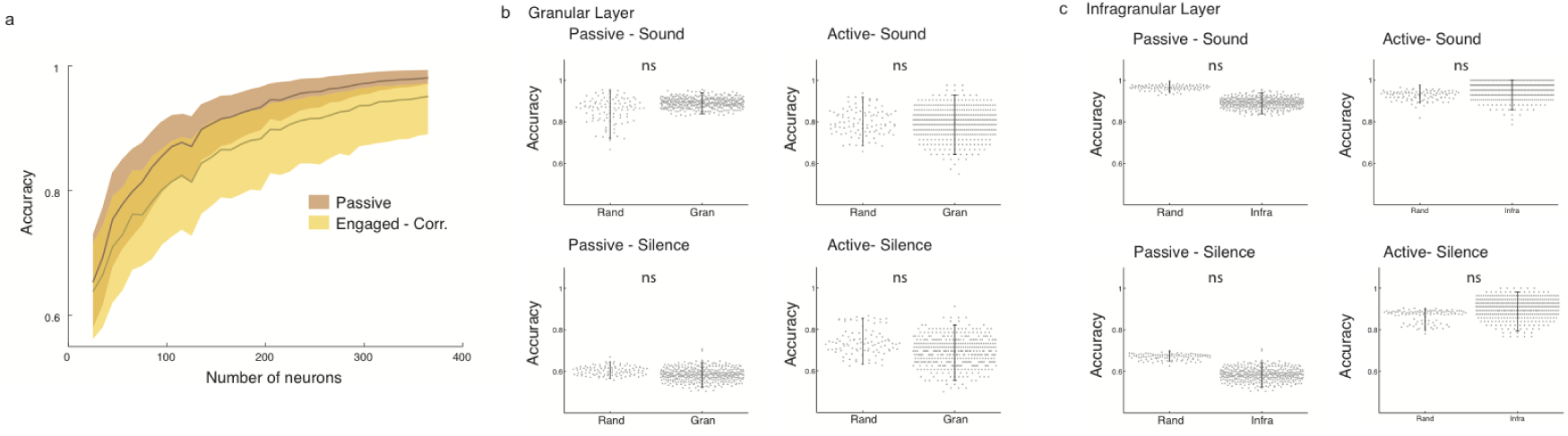
Properties of the liner classifier. *a. Effect of randomly adding units on decoding accuracy during the sound period. Error bar: 95% confidence intervals over 100 random selections of units*. *b. Units taken from the granular layer only are used for classification and accuracy is compared with the same number (89) of randomly chosen units. Error bars: 95% confidence intervals. (100 sub-sampling procedures, 400 cross validations for accuracy using granular layer units; Bonferonni corrected p-value (8 tests): 0.0063; p=0.622, p=0.933, p=0.624, p=0.618)* *c. Same as b but for infragranular layer (273 units). Error bars: 95% confidence intervals. (100 sub-sampling procedures, 400 cross validations for accuracy using granular layer units; Bonferonni corrected p-value (8 tests): 0.0063; p=0.0067, p=0.51, p=0.015, p=0.48)*

**Fig S5.**
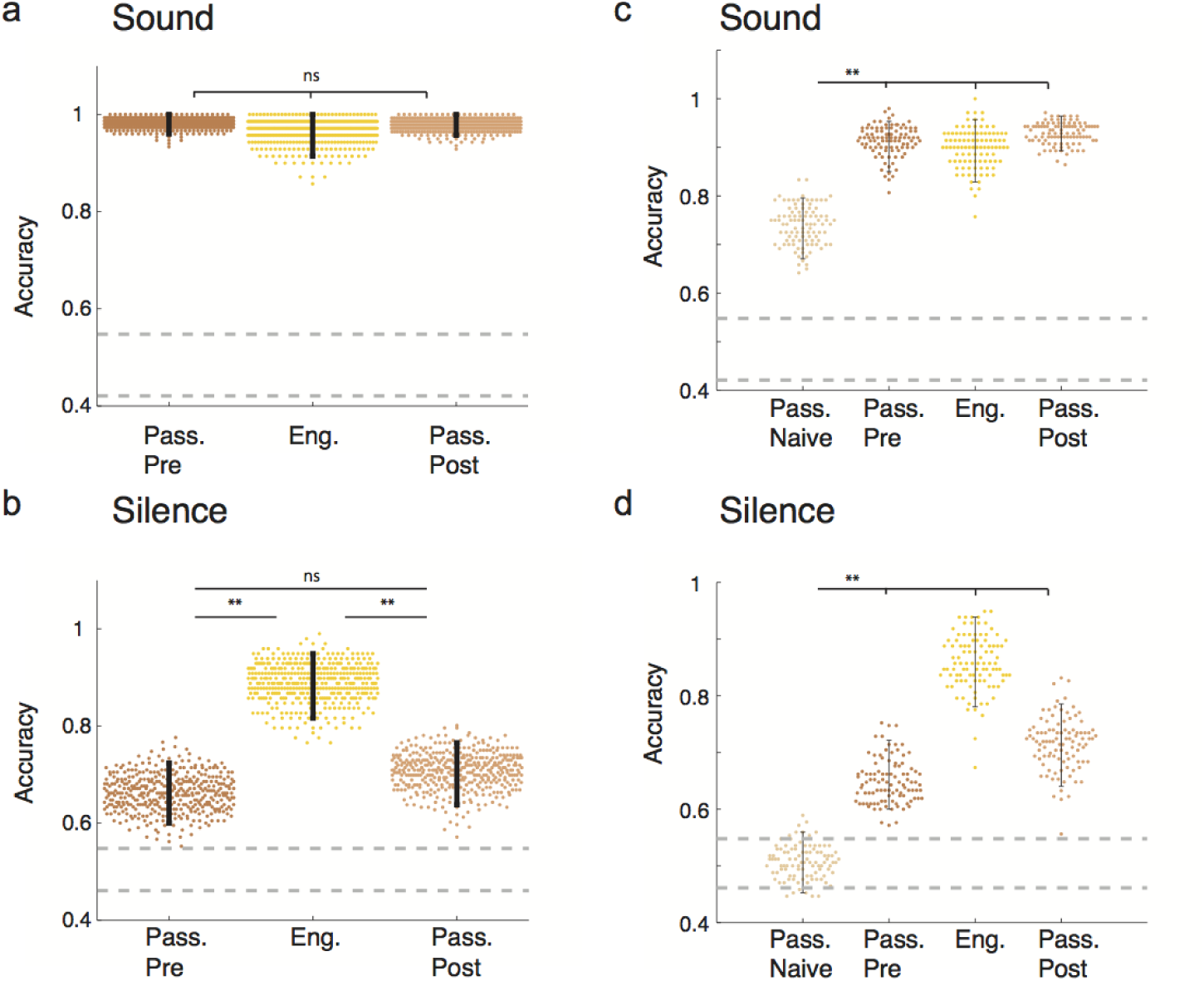
Comparison of passive session before and after behavior. *a. Comparison of accuracy during the sound period in the passive state before behavior, the task-engaged state and the passive state after behavior. Error bars represent 95% confidence intervals. (n=400 cross validations; pas.pre/eng: p=0.45, pas.pre/pas.post: p=0.74, eng/pas.post: p=0.58)*. *b. Comparison of accuracy during the silence period as in a. (n=400 cross validations; Bonferonni corrected p-value (3 tests): 0.0167; pas.pre/eng: p<0.0025, pas.pre/pas.post: p=0.43, eng/pas.post:, p<0.0025; **: p<0.01)* *c. Comparison of accuracy during the sound period in a naive animal with the passive state before behavior, the task-engaged state and the passive state after behavior in trained animals. For classification, the number of units in the trained animals was downsampled to the same number (222) as those recorded in the naive animal to allow for comparison. Error bars represent 95% confidence intervals. (n=100 cross validations after random downsampling; Bonferonni corrected p-value (3 tests): 0.0167; nve/pas.pre, nve/pas.post,nve/eng: p<0.0025;**: p<0.01)* *d. Comparison of accuracy during the silence period as in c. (n=100 cross validations after random downsampling; Bonferonni corrected p-value (3 tests): 0.0167; nve/pas.pre, nve/pas.post, nve/eng: p<0.0025;**: p<0.01)*

**Fig S6.**
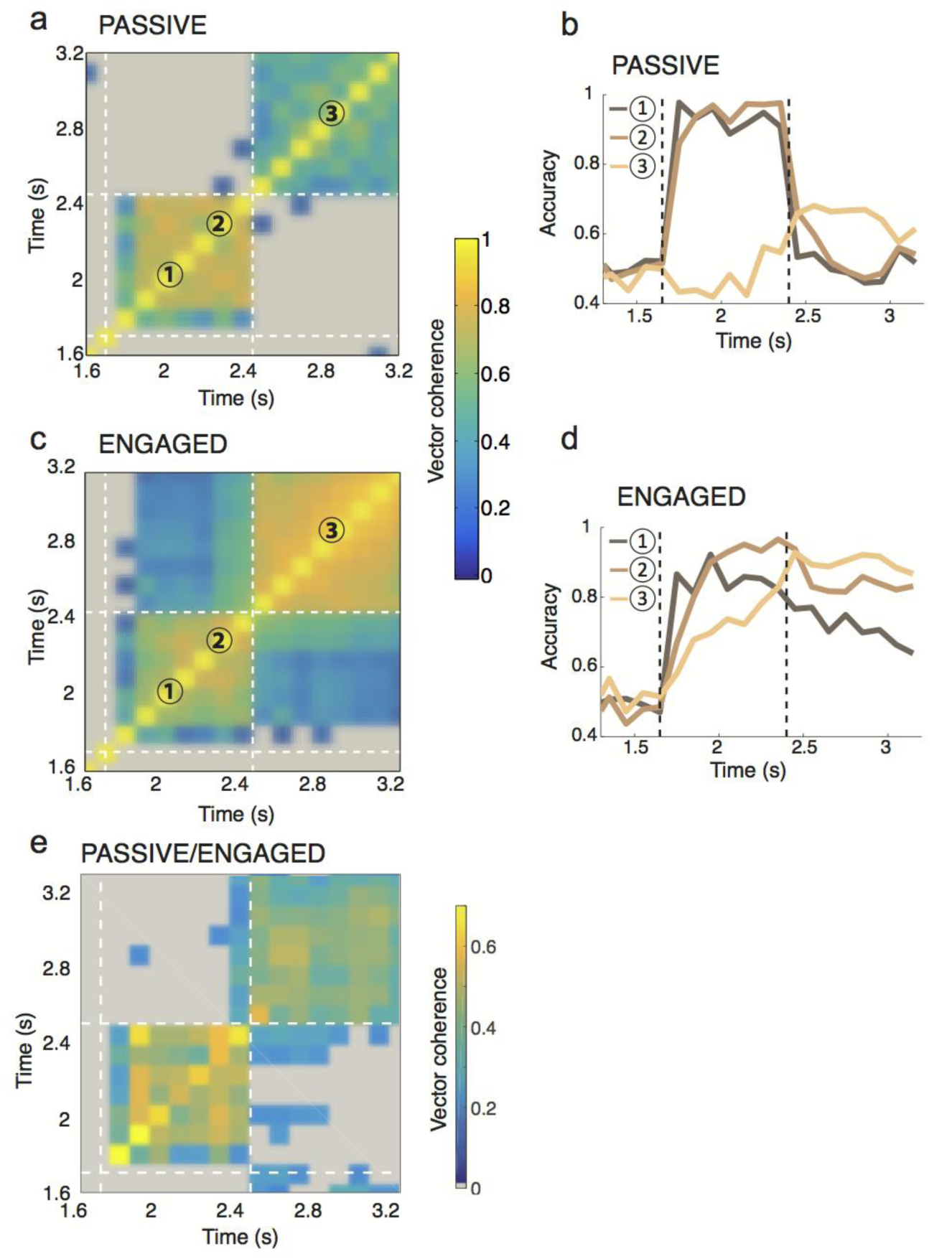
Comparison of classifiers determined at different time-points and sessions. *a. Classifier evolution in the passive state is shown in colour as the correlation between decoding vectors at one time (y-axis) versus another (x-axis). Squares with below chance correlation values are shown in grey. Here, in the passive state, coding is homogeneous throughout the sound but does not allow for significant decoding in the silent period*. *b. Decoding accuracy in the passive state using a decoder trained on the early (1) or late (2) sound or silence (3) periods. Accuracy is high throughout the sound for both early and late sound training but rapidly falls off during the silence. The decoder trained during the silence is only above chance after the sound has ended*. *c. Classifier evolution in the task-engaged state as in (a). During the silence, coding is homogeneous*. *d. As in (b) for the task-engaged state. The decoder trained during the early sound is specific to this period and performs poorly during the silence. Conversely, training late in the sound increases performance during the silence but decreases performance at the beginning of the sound. The accuracy of a decoder trained during the silence ramps up during sound presentation*. *e. Correlation of passive and engaged decoding vectors throughout the trial. Vectors show stronger similarity during the sound than the silence between states. Note the different color scale, correlation between states is as expected lower than within states*.

**Fig S7.**
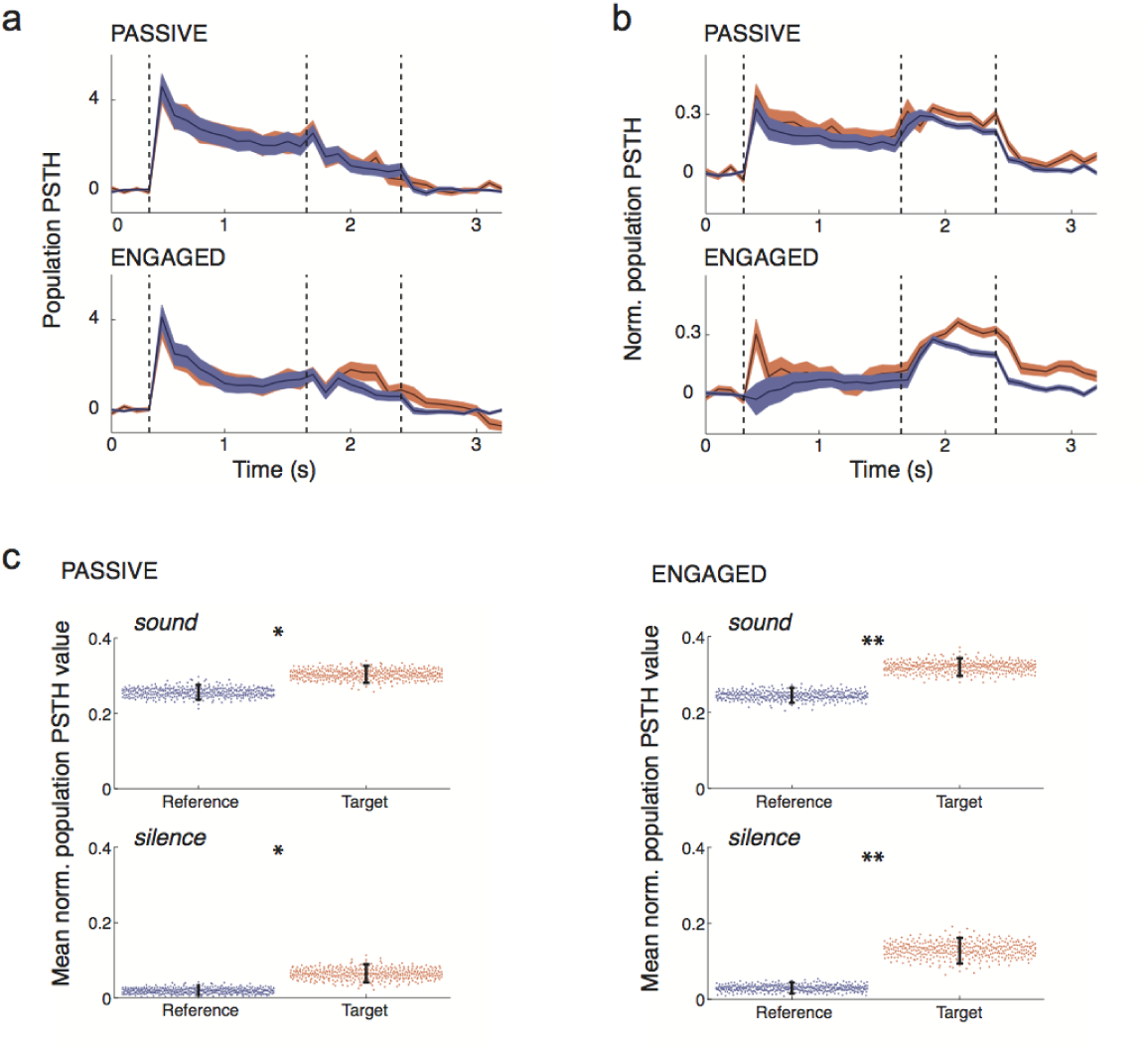
Comparing A1 population-averaged responses to target and reference stimuli. *a. Average population PSTH on reference and target trials in the passive and task-engaged states. The PSTH of each neuron is baseline subtracted and then all PSTHs are averaged. Error bars: 95% C.I. after bootstrapping 400 times over all neurons (n=370)*. *b. Average normalized population PSTH on reference and target trials in the passive and task-engaged states. The PSTH of each neuron is baseline subtracted, corrected for the sign of its peak response to reference or target and normalized to its maximal response across states and stimuli. All normalized PSTHs are then averaged. Error bars: 95% C.I. after bootstrapping 400 times over all neurons (n=370)*. *c. Distance of reference and target from baseline after normalization as in (b). Results are shown for both states during the sound or the silence period. Error bars represent 95% confidence intervals. (n=400 cross validations; pass: p=0.025 & p=0.025, eng: p<0.0025 & p<0.0025;*: p<0.05; **: p<0.01)*

**FigS8.**
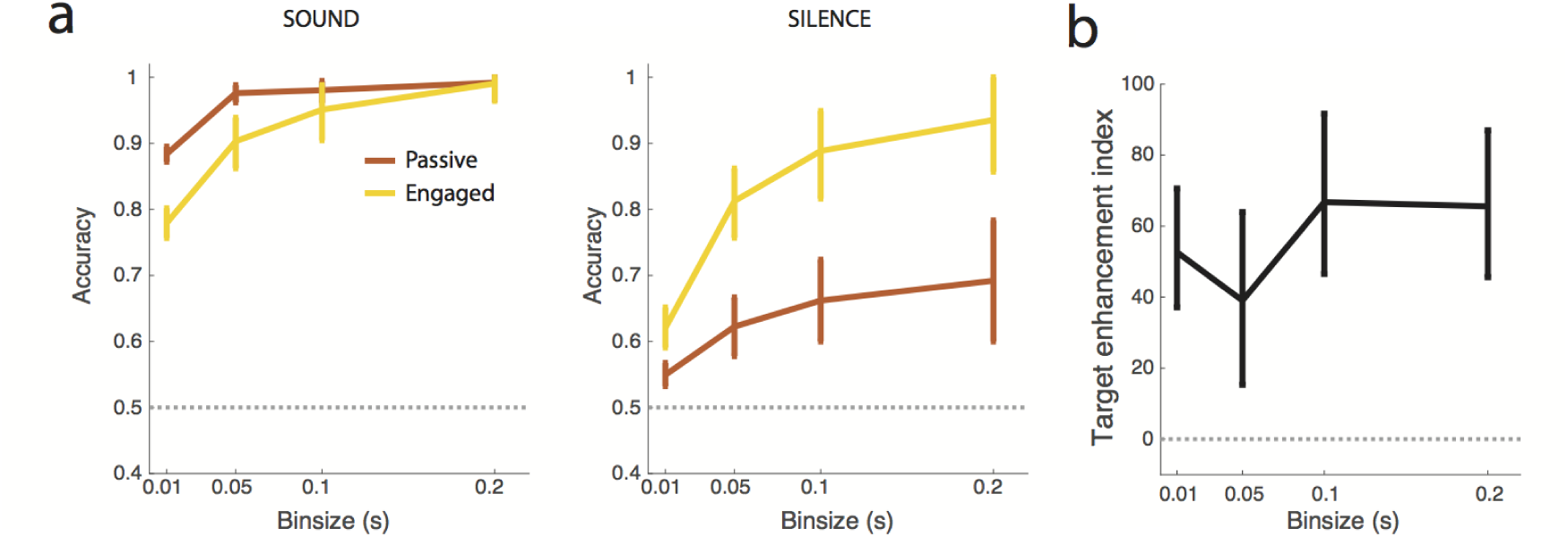
Robustness of stimulus representation characteristics across a range of time scales. *a. Accuracy of decoding during the sound (left) and silence (right) period in passive and engaged states calculated using a classifier determined with time bins of varying size. Error bars represent 95% confidence intervals. (n=400 cross validations)* *b. Index of target enhancement by task engagement calculated during the sound period using a classifier determined with time bins of varying size. Note that for all time bins the value if significantly greater than 0, indicating a systematic enhancement of target driven encoding in the engaged state. Error bars represent 95% confidence intervals. (n=400 cross validations)*

**Fig S9.**
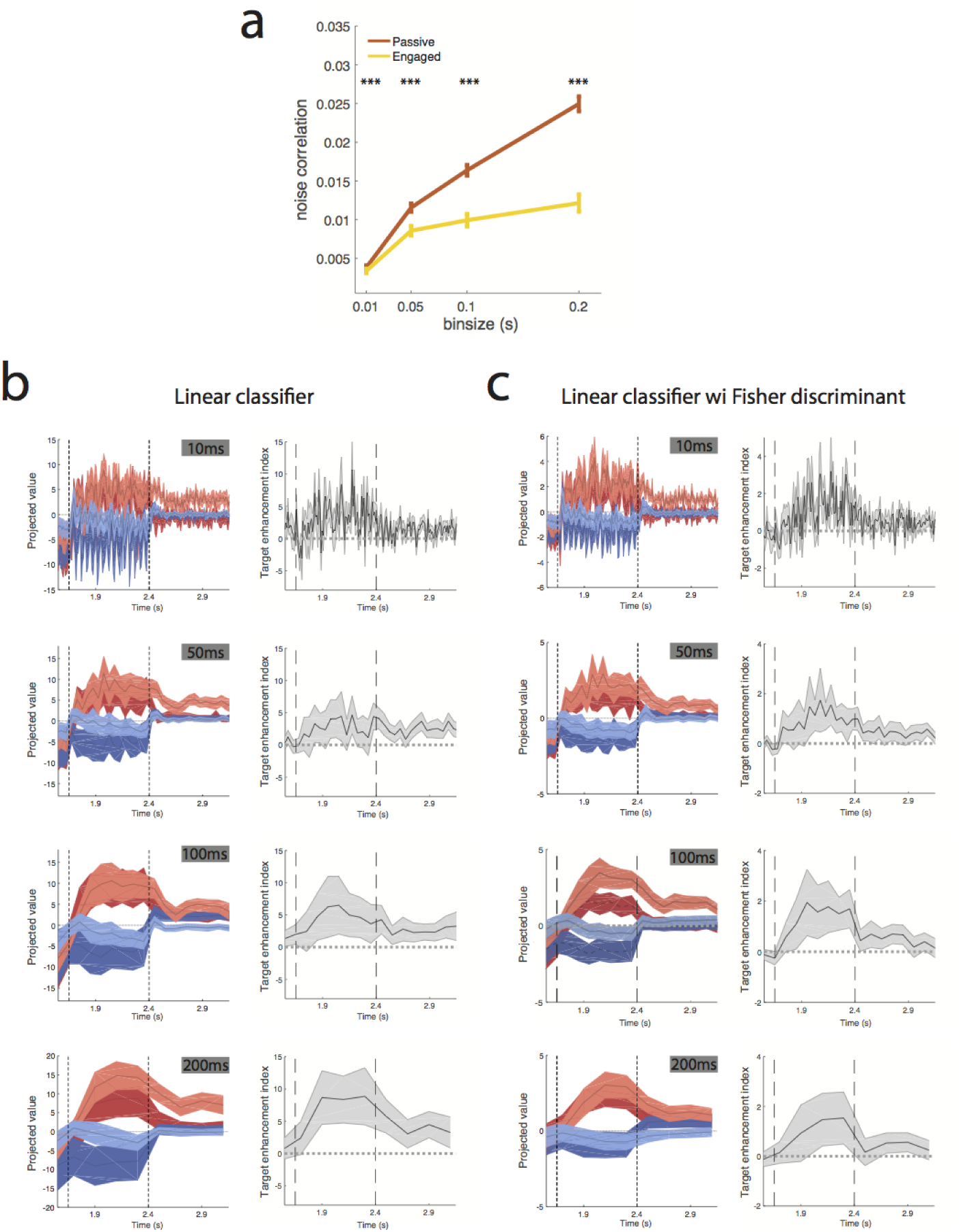
Reduced noise correlations in the engaged state does not affect enhanced asymmetry at multiple time scales. *a. Mean noise correlation in passive and engaged state using time bins of varying duration. Error bars represent s.e.m over n=3361 pairs(two-sided Wilcoxon signed rank, n=3361 pairs; zval=4.05, p=4.9E-5; zval=7.91, p=2.4E-15; zval=10.33, p=4.9E-25; zval=12.33, p=6.0E-35; ***:p<0.001)* *b. Projection onto the decoding axis determined during the sound period of trial-averaged reference (blue) and target (ref) activity during the passive (dark colors) and the active (light colors) sessions and index of target enhancement by task engagement (as in Fig5&8). Time bins of various size were used to define the decoding vector for projection. Note that for easy comparison with the Fisher discriminant analysis, decoding was done on each session individually and then the results for all sessions were averaged*. *c. As in b, for decoding vector defined using Fisher discriminant analysis*.

**Fig S10.**
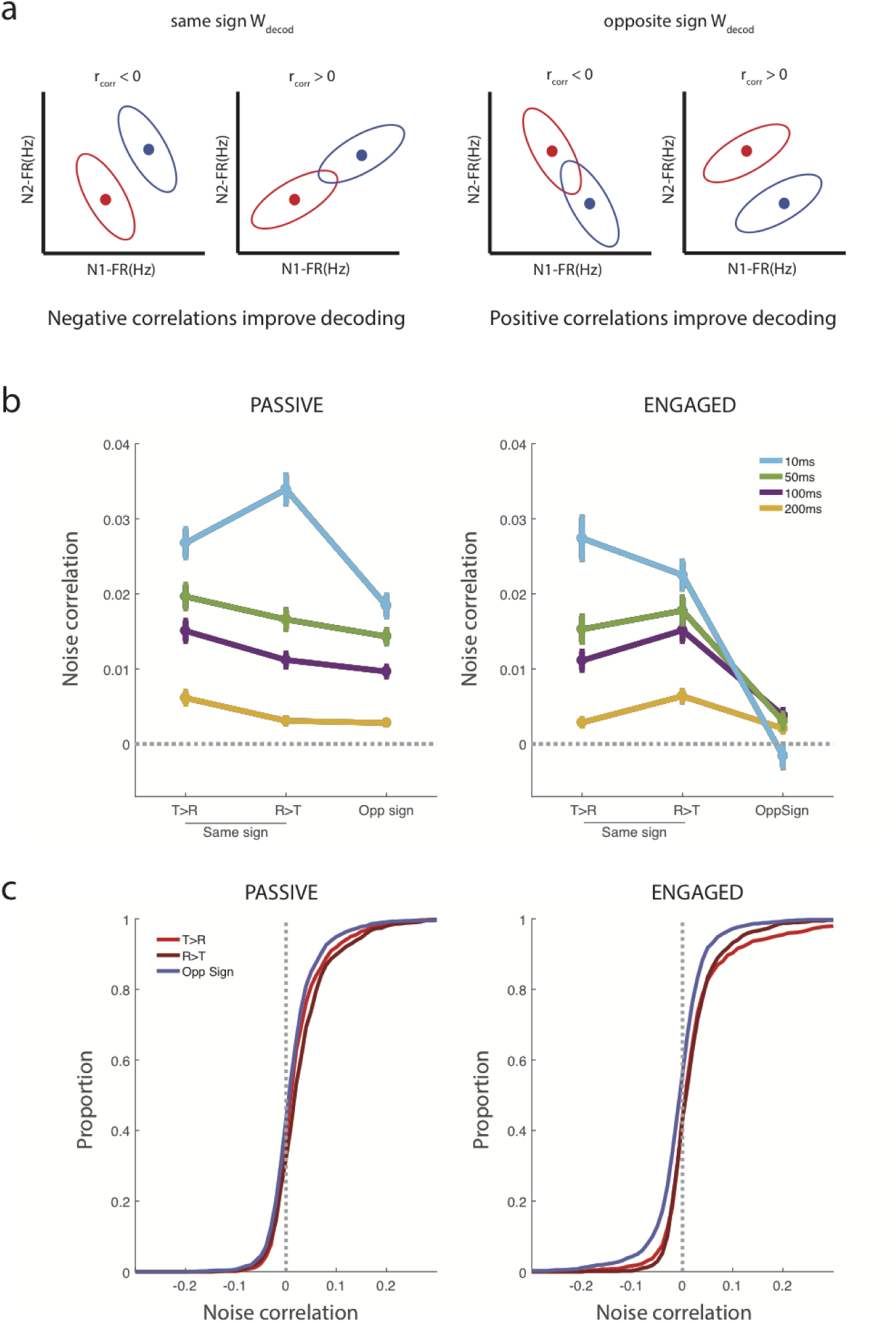
Reduction in noise correlations during taks engagement specifically impacts oppositely tuned units. *a. Schematic illustrating the relationship of ‘signal’ (decoding weight) and ‘noise’ correlations between units. Dots represent the mean target and references responses for two fictive neurons, whereas ellipses show the variance. Negative but not positive noise correlations improve stimulus discrimination for units that have the same sign of decoding weight (ie both are target-preferring or both are reference preferring) whereas the opposite if true of units with opposite sign decoding weights*. *b. Average noise correlations for units with the same or opposite sign of decoding weight in the passive (left) or engaged (right) state. In the engaged state noise correlations strongly shift towards reduced correlations for all bin sizes used in the analysis*. *c. Cumulative distribution of noise correlations for units with the same or opposite sign of decoding weight in the passive (left) or engaged (right) state. Note that the distributions are similar in the passive state whereas in the active state there is a clear shift of the noise correlations for units of opposite decoding weight sign towards lower values. There is a clear enhancement of negative correlation values*.

**Fig S11.**
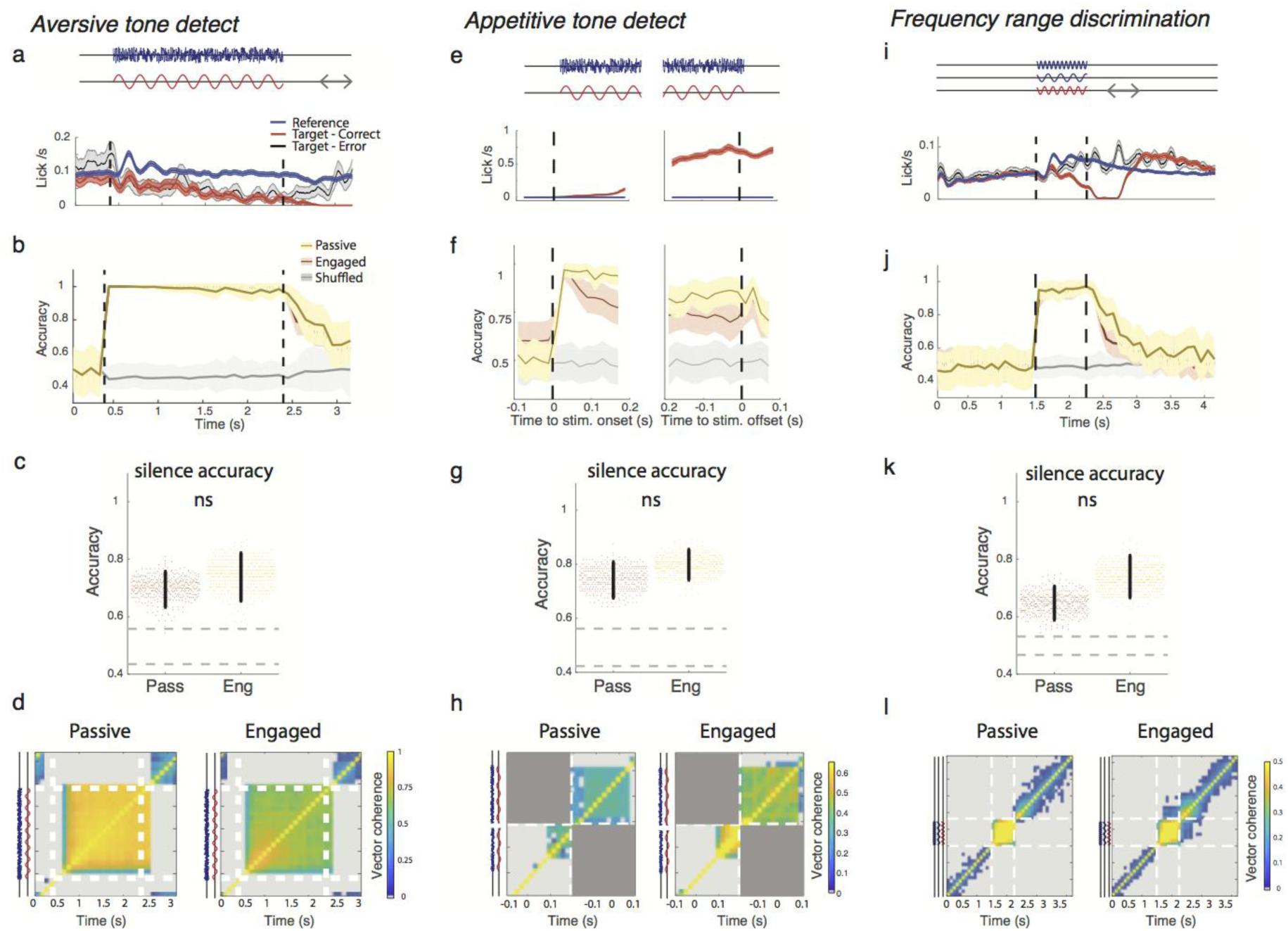
Task structure and decoding of reference/target activity in a range of auditory go/no-go tasks. *Three different tasks are considered: aversive tone detect (a-d), appetitive tone detect (e-h) and frequency range discrimination (i-l). Note that all analysis in this figure is done after excluding lick-responsive units for these tasks using the method described in Fig 4*. *a, e, i. Top: Schematic of trial structure illustrating reference and target trials. Gray arrows show response window for the aversive tasks. Bottom: Licking frequency during correct target (red), reference (blue) and target error (gray) trials. Error bars are s.e.m over all trials*. *b, f, j. Accuracy of stimulus classification in passive and engaged states. In grey, chance level performance evaluated on label-shuffled trials. Error bars represent 1 std calculated over 400 cross-validations*. *c,g,k. Mean classifier accuracy during the post-sound silence period in passive and engaged conditions. Gray dotted lines give 95% confidence interval of shuffled trials. Error bars represent 95% confidence intervals. Note that accuracy is systematically above chance level in both conditions but does not change between the passive to the engaged state. (n=400 cross validations; p=0.21,0.18,0.055)* *d,h,l. Classifier evolution in the passive (left) and engaged (right) state is shown in color as the correlation between decoding vectors at one time (y-axis) versus another (x-axis). Squares with below chance correlation values are shown in grey. For the appetitive tone detect task the overlap between sound onset and sound offset periods is not calculated as the difference in trial durations causes different overlaps in time on a trial to trial basis between the two. Note that the sound and silence periods in all tasks rely on different decoding vectors and in the case of the frequency range discrimination task, there is a progressive shift in the engaged state between decoders*.

**FigS 12.**
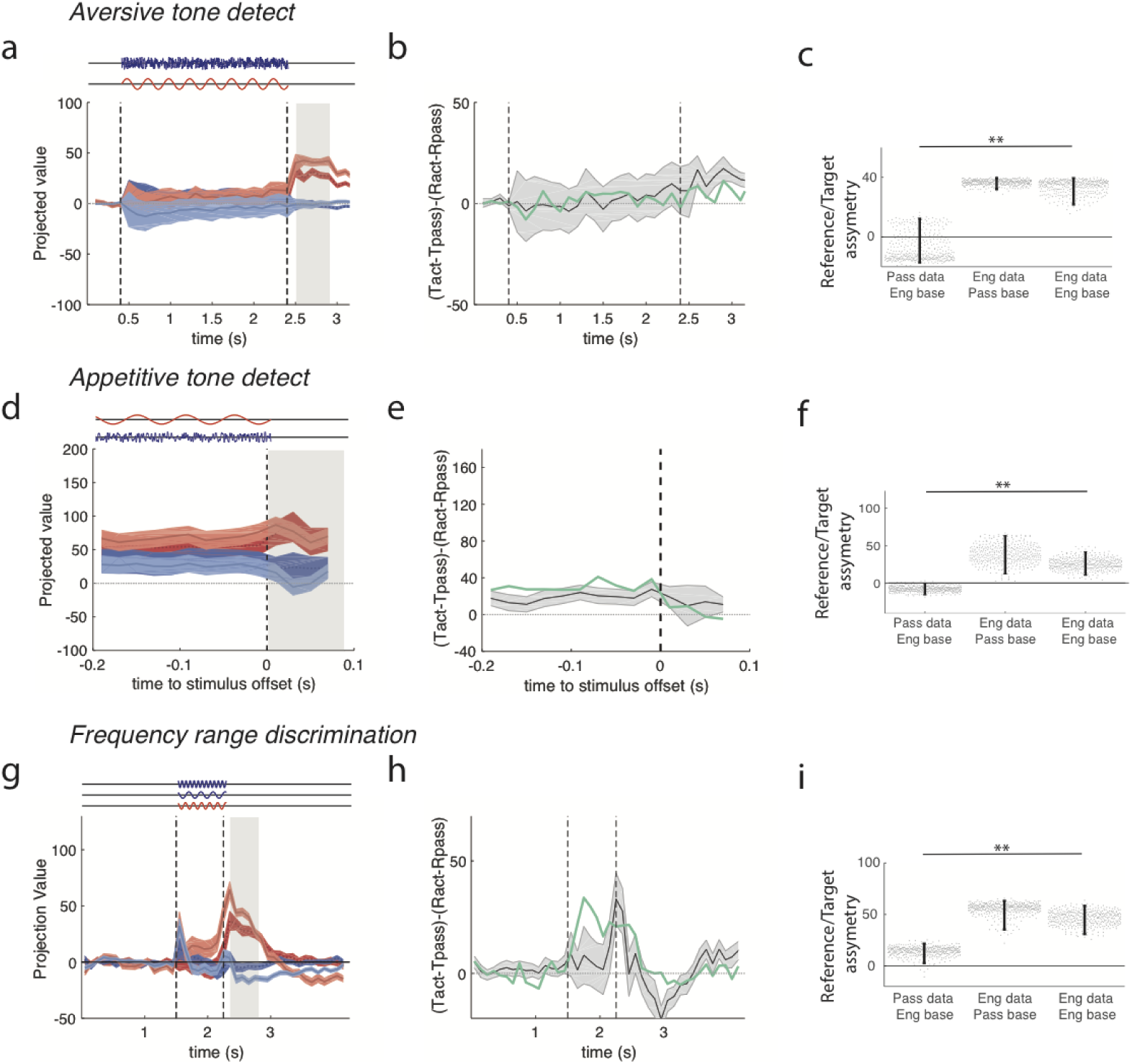
Asymmetric encoding of target and reference stimuli in a range of auditory go/no-go tasks during the post-sound silence. *a,d,g Projection of onto the decoding axis determined during the post-sound silence period of trial-averaged reference (blue) and target (ref) activity during the passive (dark colors) and the active (light colors) sessions. A baseline value computed from pre-stimulus spontaneous activity was subtracted for each neuron, so that the origin corresponds to the projection of spontaneous activity (shown by black line). Note that there is a tendency for the target-driven activity to be further from the baseline in the active state and/or the reference-driven activity to be closer. The periods used to construct the decoding axis are shaded in gray. Error bars represent 1 std calculated using decoding vectors from cross-validation (n=400)*. *b,e,h Index of target enhancement by task engagement based on projections using the decoding axis determined during post-sound silence. In green same index instead giving the same weight to all units. The difference between the green and black curved indicates that the change in asymmetry induced by task engagement cannot be detected using the population averaged firing rate alone. Error bars represent 1 std calculated using decoding vectors from cross-validation (n=400)*. *c,f,i Comparison of reference/target asymmetry for evoked responses in different states during the post-sound silence compared to different baselines given by passive or engaged spontaneous activity. Reference/target asymmetry is the difference of the distance of target and reference projected data to a given baseline. We examine three cases: (i) passive evoked responses, distances calculated relative to engaged spontaneous activity; (ii) engaged evoked responses, distances calculated relative to passive spontaneous activity; (iii) engaged evoked responses, distances calculated relative to engaged spontaneous activity. In all three cases, the engaged decoding axis was used for projections. Error bars represent 95% confidence intervals.(n=400 cross validations; Aversive Tone detect: p(col1,col3)<0.0025 & p(col2,col3)=0.92; Appetitive tone detect; p(col1,col3<0.025 & p(col2,col3)=0.94; Frequency range discrimination: p(col1,col3)<0.0025 & p(col2,col3)=0.9; **: p<0.01)*.

